# Preclinical evaluation of CDK4 phosphorylation predicts high sensitivity of malignant pleural mesotheliomas to CDK4/6 inhibition

**DOI:** 10.1101/2022.04.11.487857

**Authors:** Sabine Paternot, Eric Raspé, Clément Meiller, Maxime Tarabichi, Jean-Baptiste Assié, Frederick Libert, Myriam Remmelink, Xavier Bisteau, Patrick Pauwels, Yuna Blum, Nolwenn Le Stang, Séverine Tabone-Eglinger, Françoise Galateau-Sallé, Christophe Blanquart, Jan P. Van Meerbeeck, Thierry Berghmans, Didier Jean, Pierre P. Roger

**Author notes:** These authors contributed equally to this work (Eric Raspé, Clément Meiller). S. Paternot and P.P. Roger are co-corresponding authors. **Address for all correspondence related to this submission to:** Pierre P. Roger, IRIBHM, Université Libre de Bruxelles, Campus Erasme, 808 route de Lennik, B-1070 Brussels, Belgium.

## Abstract

Malignant pleural mesothelioma (MPM) is an aggressive cancer with limited therapeutic options. In this study, we evaluated the impact of CDK4/6 inhibition by palbociclib in a panel of 28 MPM cell lines, including 19 patient-derived cell lines, using a variety of approaches including RNA-sequencing. Palbociclib used alone sufficed to strongly and durably inhibit the proliferation of 23 MPM cell lines, indicating a unique sensitivity of MPM to CDK4/6 inhibition. Importantly, insensitivity to palbociclib was mostly explained by the lack of active T172-phosphorylated CDK4. This was associated with the high p16^INK4A^ (*CDKN2A*) levels that accompany *RB1* defects or inactivation, and also (unexpectedly) cyclin E1 over-expression in the presence of wild-type *RB1*. Prolonged treatment with palbociclib irreversibly inhibited proliferation despite re-induction of cell cycle genes upon drug washout. A senescence-associated secretory phenotype including various potentially immunogenic components was also irreversibly induced. Phosphorylated CDK4 was detected in 80% of 47 MPM tumors indicating their intrinsic sensitivity to CDK4/6 inhibitors. The absence of this phosphorylation in some highly proliferative MPM tumors was linked to partial deletions of *RB1*, leading to very high p16 (*CDKN2A*) expression. Our study strongly supports the clinical evaluation of CDK4/6 inhibitory drugs for MPM treatment, in monotherapy or combination therapy.

## Introduction

Malignant pleural mesothelioma (MPM) is a rare and aggressive tumor mostly associated with asbestos exposure. Despite the ban of asbestos in several countries, it remains a major public health concern worldwide. MPMs are classified into three main histological subtypes with different prognosis: epithelioid, biphasic and sarcomatoid (1). As current treatment options (cisplatin+pemetrexed combined with bevacizumab if available) (2–4) only extend survival for a few months, alternative therapies with predictive markers based on the MPM biology are desirable (5–8). Recently, dual immune checkpoint inhibition with nivolumab plus ipilimumab has demonstrated superiority to platinum plus pemetrexed in MPMs with non-epithelioid histologies (9), but this remains questioned for epithelioid MPMs (4).

MPMs are mostly characterized by frequent deletions (50-80%) of the *CDKN2A/B* locus (encoding the CDK4/6 inhibitors p16^INK4A^ and p15^INK4B^) (6, 10–12), which are associated with shorter overall survival (13). Other recurrent defects include mutations affecting the neurofibromatosis 2 (*NF2*) tumor suppressor encoding merlin and the BRCA1 associated protein 1 (*BAP1).* NF2, as a key player in the Hippo pathway involved in cell contact growth inhibition (10), negatively regulates the expression of cyclin D1 (14). These different defects are expected to deregulate the activity of CDK4 and cause the observed hyper-proliferation. On the other hand, the onco-suppressor pRb (*RB1*), the main target of CDK4, has been found to be intact in most MPMs, pointing out these tumors as paradigmatic targets for CDK4/6 inhibitors (6, 12). However, a recent study reported *RB1* deletions (possibly mostly monoallelic) in 26% of 118 MPM samples including in one third of tumors with *CDKN2A* deletions (15).

Pharmacological inhibition of CDK4/6 has emerged as a promising anti-cancer strategy. Palbociclib, ribociclib and abemaciclib, combined with endocrine therapy, recently became a standard-of-care as first-line treatments for advanced ER-positive breast cancers. Despite mild side effects (cytopenia,..), these CDK4/6 inhibitors (CDK4/6i) induce an ‘unprecedented improvement of progression-free survival’ (16) and also significantly improve the overall survival (17, 18). In different cancer models, specific inhibition of CDK4/6 not only induces the cell cycle arrest but also a senescent-like state (19–23) including a senescence-associated secretory phenotype (SASP) (21, 24). However, because CDK4/6 inhibition is often insufficient to fully control the tumor progression, rationally developed doublet or triplet combination therapies of CDK4/6i are evaluated in various cancers, for instance with kinase inhibitors (e.g. MEK, PI3K, mTOR, EGFR inhibitors,...) (21), genotoxic therapies (25, 26) and immunotherapies (21, 25, 27, 28).

Cyclin D-CDK4/6 are the first CDK complexes to be activated during the G1 phase in response to mitogenic/oncogenic stimuli (29–31). Their main function consists in the inactivation of the pRb (*RB1*) anti-oncogene (32). Insensitivity to CDK4/6i is generally ascribed to the functional loss or inactivation of pRb (33). However, CDK4 and CDK6 might also control the cell cycle progression by phosphorylating a growing list of other proteins including p107, p130, FoxM1, Smad3 (31, 34–38). pRb inactivating phosphorylations in committed cells are maintained by a positive feedback loop linking pRb to E2F-dependent transcription of *CCNE1* (cyclin E1), which activates CDK2, further enhancing the inactivation of pRb (39). CDK4 activity requires its binding to a cyclin D (*CCND1-3* genes) competed by INK4 CDK4 inhibitors like p16 (*CDKN2A-D* genes) (29, 40). To be active, CDK4 also needs to be phosphorylated on T172 (41). Our group has identified the activating T172-phosphorylation of cyclin D-bound CDK4 as the last distinctly regulated step in CDK4 activation, determining the cell cycle commitment in pRb-proficient cells (30, 42–47). In breast cancer tumors and cell lines, we have reported that the highly variable abundance of phospho-CDK4 signals the presence or absence of active CDK4 targeted by CDK4/6i, which was associated with the sensitivity or insensitivity of tumor cells to palbociclib (48). CDK4 T172-phosphorylation might thus be the best biomarker of potential tumor sensitivity to CDK4/6i.

Discordant results were reported in the few studies evaluating CDK4/6i in MPM cell lines (15, 49, 50). Nevertheless, a single-arm, open-label, phase II trial of 27 MPM patients recently showed a promising clinical activity of abemaciclib (51). Here, we demonstrate and characterize the unique responsiveness of most MPM cell lines to CDK4/6 inhibition. In a few cell lines, a complete insensitivity to palbociclib is nevertheless observed. It is mostly associated with the absence of phosphorylated CDK4, which can occur even in the absence of *RB1* defect. CDK4 phosphorylation is detected in about 80% of MPM tumors, which are therefore predicted to be responsive to CDK4/6i. CDK4 phosphorylation is nevertheless undetectable in a subset of MPMs, which should be considered in future clinical settings involving CDK4/6i.

## Results

### Most MPM cell lines exhibit a high sensitivity to CDK4/6 inhibition, which correlates with the presence of phosphorylated CDK4

We selected a panel of well-characterized MPM cell lines representative of the three histotypes and comprising both commercial and patient-derived cell lines including low-passage ones (Supplementary Table S1). MPP89 was chosen because it harbors a mutated *RB1*. Normal mesothelial cells immortalized by SV40 (MeT-5A) were also used as a model of pRb-deficient cells. Treatment with palbociclib for 24 h prevented the cell cycle progression in a concentration-dependent manner in 23 of 28 cell lines (Fig. 1A, Supplementary Fig. S1). In the majority of these, 1 μM palbociclib reproducibly reduced the number of DNA replicating cells by more than 90% (Fig. 1A,B). The effect of CDK4/6 inhibition was less pronounced when measured using MTT or sulforhodamine viability assays, confirming the cytostatic effect of palbociclib (Supplementary Fig. S1). Similar results were obtained with ribociclib and abemaciclib (Supplementary Fig. S1A,B).

**Figure 1.**
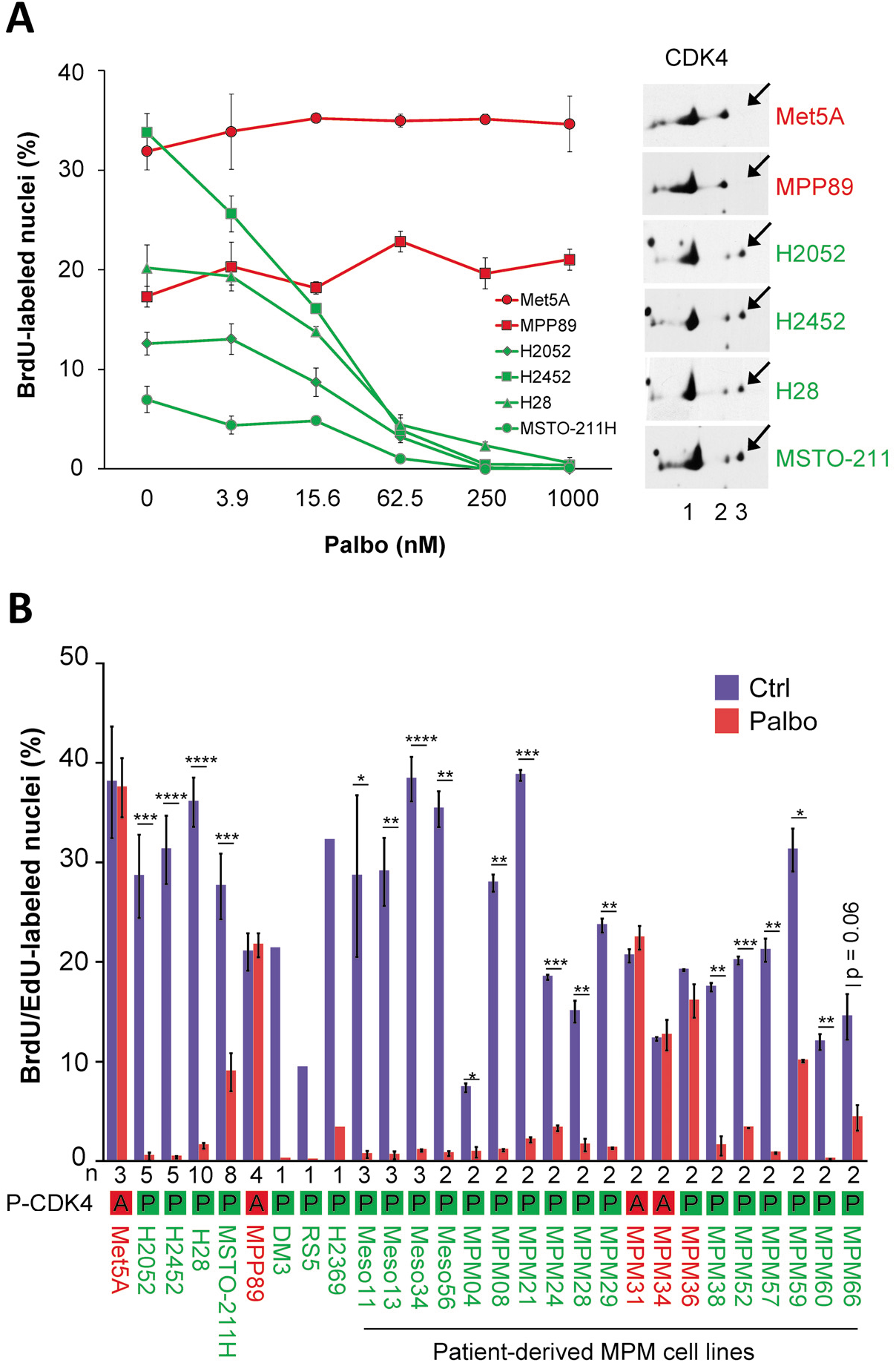
Sensitivity of MPM cell lines to palbociclib is associated with CDK4 phosphorylation. Sensitive cell lines are in green, resistant cell lines are in red. **A** Left panel. DNA synthesis in MPM cell lines treated for 24 h with increasing concentrations of palbociclib (Palbo). Data show one representative experiment (mean +/− SD of duplicated dishes). Right panel. CDK4 immunodetection after separation by 2D-gel electrophoresis. Arrows indicate the position of the T172-phosphorylated form of CDK4 (spot 3). **B** DNA synthesis in cells treated for 24 h with DMSO (Ctrl) or palbociclib (Palbo) at 1 μM. Data represent mean +/− SEM of n independent experiments, as mentioned below the graph. P-CDK4 indicates the phosphorylation status of CDK4: A, absent and P, phosphorylated. *p<0.05, **p<0.01, ***p<0.001, ****p<0.0001 (Student’s t-test).

In parallel, we evaluated in these cell lines the presence of the activated form of CDK4 phosphorylated on T172. This was achieved by 2D-gel electrophoresis separation of CDK4 forms as previously characterized (42, 48) (Fig. 1A, Supplementary Fig. S1). Whereas the T172-phosphorylated form of CDK4 was detected in all the palbociclib-sensitive cell lines, it was absent in four of five resistant ones (MPP89, MeT-5A, MPM_31, MPM_34; Fig. 1B, Supplementary Fig. S1). With the noticeable exception of resistant MPM_36 cells that displayed a low level of CDK4 phosphorylation (Supplementary Fig. S1), the presence or absence of CDK4 T172-phosphorylation thus correctly predicted the sensitivity or insensitivity to CDK4/6 inhibition in MPM cell lines, similarly to our initial observations in breast cancer cell lines (48).

### Absence of CDK4 phosphorylation in resistant cells is due to high p16 levels associated or not with pRb defect

To better understand the mechanisms involved in palbociclib resistance and in the lack of CDK4 phosphorylation, the cell lines were characterized by immunodetection of key cell cycle related proteins and by RNA-sequencing (RNA-seq) for 19 of them. Insensitive cell lines lacking phospho-CDK4 were distinguished by strongly elevated p16 levels, consistent with high *CDKN2A* mRNA expressions (Fig. 2A,B; Supplementary Fig. S2; Supplementary Table S1). In that situation, coimmunoprecipitation experiments showed that p16 prevents the activating phosphorylation of CDK4 by impairing its binding to D-type cyclins (Fig. 2C). Constitutive loss of function of pRb, the most obvious driver of resistance to CDK4/6i, is known to lead to elevated *CDKN2A* mRNA levels (52, 53). Inactivation of pRb by SV40 in MeT-5A and its mutation in MPP89 could thus explain the high levels of p16 in these two cell lines and the resulting impairment of CDK4 phosphorylation. This role of p16 was further demonstrated by the specific case of MPM_36 cells, which harbors both *RB1* and *CDKN2A* biallelic deletions (Fig. 2A, Supplementary Table S1). Indeed, MPM_36 was the only pRb-deficient palbociclib-resistant cell line that presented a detectable phosphorylation of CDK4; reproducing the exceptional situation observed in DU4475 breast cancer cells (48).

By contrast, MPM_31 and MPM_34 presented high *CDKN2A* mRNA and p16 levels despite normal expression and phosphorylation of pRb (Fig. 2A,B, Supplementary Fig. S2). No *RB1* defect (mutation or deletion) was found by RNA-seq covering all *RB1* exons of the MPM_31 and MPM_34 resistant cells (Supplementary Table S1). Inactivation of pRb by SV40 large T was also excluded by RNA-seq analysis (Supplementary Table S2). On the other hand, these cells were among those expressing the highest abundances of cyclin E1 at the protein and mRNA levels (Fig. 2A,B, Supplementary Fig. S2A), consistent with the amplification of the *CCNE1* gene locus detected by SNP array (Fig. 2D). In addition, MPM_31 displayed an abnormal electrophoretic migration of p27 (two bands; Fig. 2A, Supplementary Fig. S2A), which we found to be associated with a I119T mutation of one *CDKN1B* allele that was previously reported in other cancers (Supplementary Table S1)(54–56). We also observed an abnormal electrophoretic migration of cyclin E1 in MPM_31 (Fig. 2A, Supplementary Fig. S2A), which most likely indicated a post-translational modification because it could not be explained by any genetic alteration detected by RNA-seq of *CCNE1.* To evaluate the effect of these peculiarities, we analyzed by co-immunoprecipitation the formation and activity of different CDK complexes in MPM_31 and MPM_34 as compared to three palbociclib-sensitive primary MPM cell lines (Fig. 2E,F). Similarly to MPP89 and MeT-5A, CDK4 was bound to p16 but neither to cyclin D1 nor cyclin D3 in MPM_31 and MPM_34 (Fig. 2E). A much higher pRb-kinase activity was associated with CDK2 and cyclin E1 in these two resistant cell lines (Fig. 2F). Interestingly, an increased pRb-kinase activity was also associated with p27 in MPM_31 cells. This might be ascribed to p27-bound CDK2 complexes because no association of p27 with CDK4 or CDK6 was detected in this cell line (Fig. 2F), consistent with the main association of CDK4 with p16 (Fig. 2E). As analyzed by 2D-gel electrophoresis of CDK2 (57), the relative proportion of the active form of CDK2 phosphorylated on T160, but not on Y15 or T14, was also more elevated in MPM_31 cells (Fig. 2G), suggesting a particularly elevated cdc25 phosphatase activity. Overall, these different observations suggest (for the first time to our knowledge) that a constitutive over-activation of cyclin E1-CDK2 and the resulting inactivation of pRb by phosphorylation can also lead to high p16 expression and inactivation of CDK4, explaining the resistance to CDK4/6i.

The detection of T172-phosphorylated CDK4 was a more consistent biomarker of sensitivity to palbociclib than the expression of any other key cell cycle regulatory protein. Indeed, pRb was detected not only in all the palbociclib-sensitive MPM cell lines but also in 4 of 5 resistant ones, though at much reduced levels in the *RB1*-mutated MPP89 cells (Fig. 2A). Cyclin D1 was less abundant in all the insensitive *RB1*-deficient cells, but it was present in MPM_31 and MPM_34 cells. p16 was highly expressed in 4 of 5 insensitive cell lines, but it was also present at moderate levels in the DM-3 and RS-5 sensitive cell lines. By contrast, p16 was undetectable in 21 of 23 sensitive cell lines, mostly due to deletion of *CDKN2A* or its mutation (paradoxically associated with elevated *CDKN2A* RNA levels in MPM_21 cells) (Fig. 2A,B; Supplementary Table S1). In MPM_59, loss of p16 expression might be due to methylation of the *CDKN2A* locus or mutation in its promoter region, since its RNA levels were low while no deletion or mutation were detected. It should be noted that MPM cell lines expressing normal levels of wild-type p16 grew very slowly, precluding any detailed evaluation. Two such patient-derived cell lines had to be excluded from our study because of insufficient EdU incorporation.

In brief, our results show that the cell cycle resistance to palbociclib was associated not only to pRb defects but also to the absence of CDK4 phosphorylation due to high p16 levels, even in the presence of normal pRb.

**Figure 2.**
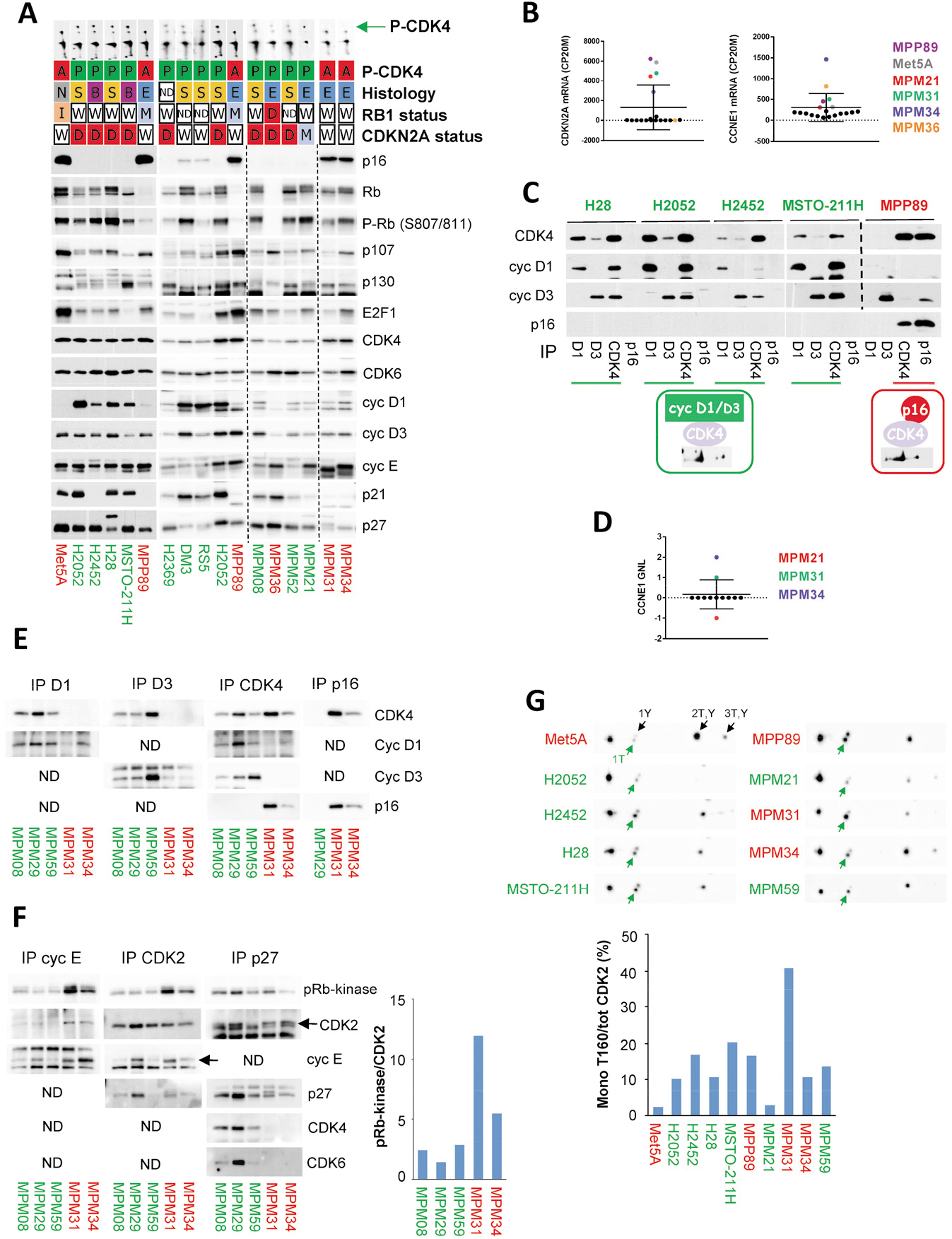
Absence of CDK4 phosphorylation in resistant cells is due to high p16 levels associated with pRb defect or high CDK2 activity. Sensitive cell lines are in green, resistant ones are in red. **A** Western Blot analysis of the indicated proteins and profile of CDK4 separated by 2D-gel electrophoresis (the green arrow indicates the position of the phosphorylated form of CDK4). Vertical dashed lines separate parts of the same blot that were assembled. The following data are given for each cell line: CDK4 phosphorylation profile (P-CDK4, A: absent, P: phosphorylated), histological subtype (E: epithelioid, S: sarcomatoid, B: biphasic, N: normal, ND: not determined), status of *RB1* and *CDKN2A* genomic locus (W: wild-type, I: wild-type but inactivated by SV40, D: deleted, M: mutated, ND: not determined). **B** *CDKN2A* and *CCNE1* mRNA expression extracted from RNA-seq data and normalized relative to the library size in counts per 20 million reads (CP20M). N= 19 cell lines. Mean of 2-3 experiments except for MPM_36 and MPM_66 (n=1). **C** Immunodetection of the indicated proteins from CDK4 complexes co-immunoprecipitated (IP) using anti-cyclin D1, cyclin D3, CDK4 or p16 antibodies. **D** Gain/normal/loss (GNL) score of *CCNE1* locus analyzed by SNP array. N = 12 cell lines. **E** Immunodetection of the indicated proteins from CDK4 complexes co-immunoprecipitated (IP) using anti-cyclin D1, cyclin D3, CDK4 or p16 antibodies. ND: not done (less relevant). **F** Co-immunoprecipitation (IP) of CDK2 complexes using anti-cyclin E1, CDK2 or p27 antibodies followed by pRb-kinase assay and immunodetection of the indicated proteins. ND: not done (less relevant). Quantification of phospho-pRb signal normalized to CDK2 is shown on the right. **G** CDK2 immunodetection after separation by 2D-gel electrophoresis. Arrows indicate the various phosphorylated forms of CDK2. 1T (active CDK2 in green): mono-T160 phosphorylation, 1Y: mono-Y15 phosphorylation, 2T,Y: double phosphorylation on T160 and Y15 and 3T,Y: triple phosphorylation on T160, Y15 and T14. Quantification of the proportion of active CDK2 phosphorylated only at T160 (mono-T160) over total CDK2 is displayed below the detections.

### Prolonged palbociclib treatment induces senescence in MPM cell lines

The effect of palbociclib on cell cycle progression was maintained for at least two weeks (Fig. 3A), contrary to the rapidly developing resistance to CDK4/6 inhibition described in breast cancer cells (58). In sensitive MPM cells, prolonged treatment (9 days) with palbociclib induced a senescent phenotype characterized by an enlarged and flat cell morphology and by increased senescent-associated β-galactosidase activity (SA-β-gal) (Fig. 3B). Moreover, this long-term pretreatment with palbociclib increased the apoptosis induced by senolytic drugs targeting Bcl-2 and Bcl-xL (59) (Fig. 3C).

**Figure 3.**
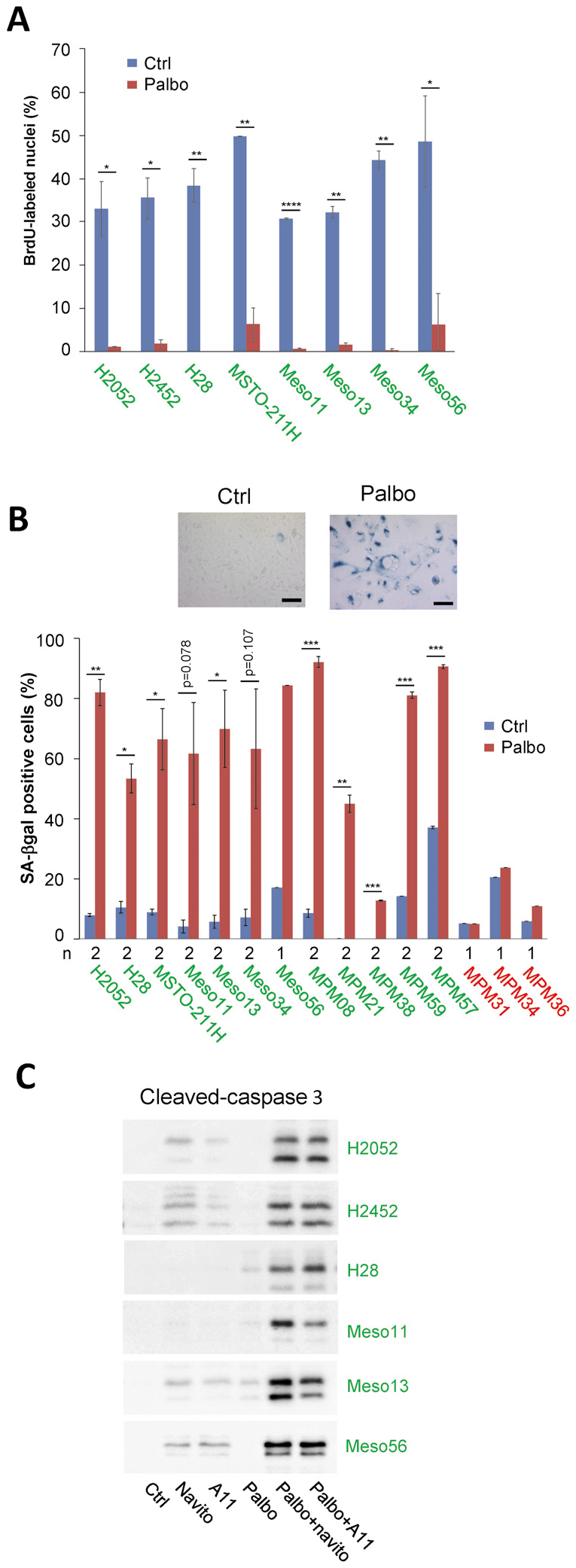
Prolonged palbociclib treatment induces senescence in MPM cell lines. **A** DNA synthesis in cells treated for 2 weeks with DMSO (Ctrl) or palbociclib (Palbo) at 1 μM. Data are presented as mean +/− SEM of 2 independent experiments. **B** Representative SA-β-gal staining of a sensitive cell line (H2052) treated for 9 days with DMSO (Ctrl) or 1 μM palbociclib (Palbo). Quantification represents the percentage of cells positive for SA-β-gal staining (mean +/− SEM of 2 independent experiments except for resistant cells and Meso56, as indicated below the graph). Scale bar, 100 μm. **C** Immunodetection of cleaved-caspase 3 in cells grown for 3 days in the absence (Ctrl) or presence of 1 μM navitoclax (navito) or A1155463 (A11) and pre-treated or not (Ctrl) for 9 days with 1 μM palbociclib (Palbo). *p<0.05, **p<0.01, ***p<0.001, ****p<0.0001 (Student’s t-test).

To further define the response of MPM cell lines to CDK4/6 inhibition, we compared the transcriptomic profiles of sensitive and resistant cell lines treated for 9-10 days with 1 μM palbociclib or DMSO (Fig. 4, Supplementary Fig. S3). RNA-seq data of selected targets were validated at the protein level by Western Blotting and the increased secretion of various components of the SASP was confirmed using cytokines arrays (Supplementary Fig. S4 and S5). Remarkably, no significant gene expression changes were observed in the insensitive cell lines (Fig. 4A,B), confirming the high specificity of this inhibitor used at 1 μM. This result also intriguingly indicated that both the absence of CDK4 phosphorylation in the presence of functional pRb (MPM_31 and MPM_34), and the loss of pRb in the presence of phosphorylated CDK4 (MPM_36), suffice to generate a complete insensitivity to palbociclib. In most sensitive cells, treatment with palbociclib induced a strong down-regulation of genes (Fig. 4C; Supplementary Fig. S3A) and proteins (Supplementary Fig. S4A,B and S5A,B) involved in cell cycle progression and DNA repair. Moreover, the long term treatment with the CDK4/6i increased the expression of many genes, the number of which even exceeded the number of down-regulated ones (Fig. 4A,B; Supplementary Tables S3 and S4). These included various genes and proteins classically involved in senescence (Fig. 4C; Supplementary Fig. S3A, S4 and S5), such as D-type cyclins, cell adhesion molecules (ICAM1, VCAM1, L1CAM), and SASP components, such as SERPINs, tissue plasminogen activator (PLAT), fibronectin (*FN1),* IGFBPs, complement C3 and VEGFA. Several pro-inflammatory cytokines (IL1A/B, IL6) and chemokines (CCL2, CCL5, CXCL8 (IL8)), as well as Major Histocompatibility complex class I, were also up-regulated.

Gene Set Enrichment Analysis identified “E2F targets” and “DNA repair” among the top down-regulated “Hallmarks” pathways in response to CDK4/6 inhibition (Fig. 4D,E; Supplementary Fig. S3B; Supplementary Tables S5 and S6). Related “Kegg” pathways including “Cell cycle” and “Base excision repair” were also strongly down-regulated by palbociclib. On the other hand, signatures related to senescence and immune response such as “Fridman senescence up”, “TNFA signaling via NFKB”, “Cytokine-cytokine receptor interaction”, “Natural killer cell-mediated cytotoxicity”, “JAK STAT signaling pathway”, “Antigen processing and presentation”, “Leukocyte transendothelial migration” were enriched in palbociclib-treated cells (Fig. 4D,E; Supplementary Fig. S3B; Supplementary Tables S5 and S6). “Interferon gamma response” and “Interferon alpha response” were also very significantly up-regulated by palbociclib, but interestingly were not associated to a detectable expression of any type I, II or III interferon genes, consistent with the loss of their locus or low basal expression, as reported previously (60). “Complement”, “Coagulation”, “Angiogenesis” and “Apoptosis” were also increased by palbociclib treatment. Of note, “Epithelial Mesenchymal Transition” was the top up-regulated “Hallmark” pathway (Fig. 4D,E). However, this most likely reflected the profound remodeling of the extracellular matrix and cytoskeleton that is also associated with senescence. Indeed, palbociclib up-regulated both “epithelial” and “mesenchymal” markers (defined as in ref.(61)) (Fig. 4E). Moreover, classical changes linked to EMT such as reduced E-cadherin expression and up-regulation of vimentin, N-cadherin and several EMT transcriptional inducers were not observed (Supplementary Fig.S3A). Analysis of enriched pathways in individual cell lines illustrated some heterogeneity in the pathways up-regulated by palbociclib. It also highlighted an opposite regulation of the interferon signaling, in particular in MPM_08 cells that were characterized by high basal expression of *STAT1* and of interferon-induced genes (*ISGs*) (Fig. 4E; Supplementary Table S6). Most importantly, the inhibition of CDK4/6 was nevertheless able to reverse, in all the evaluated MPM cell lines, the “immune resistance program” that promotes T cell exclusion and resistance to immunotherapy in melanoma (62) (Fig. 4D,E).

**Figure 4.**
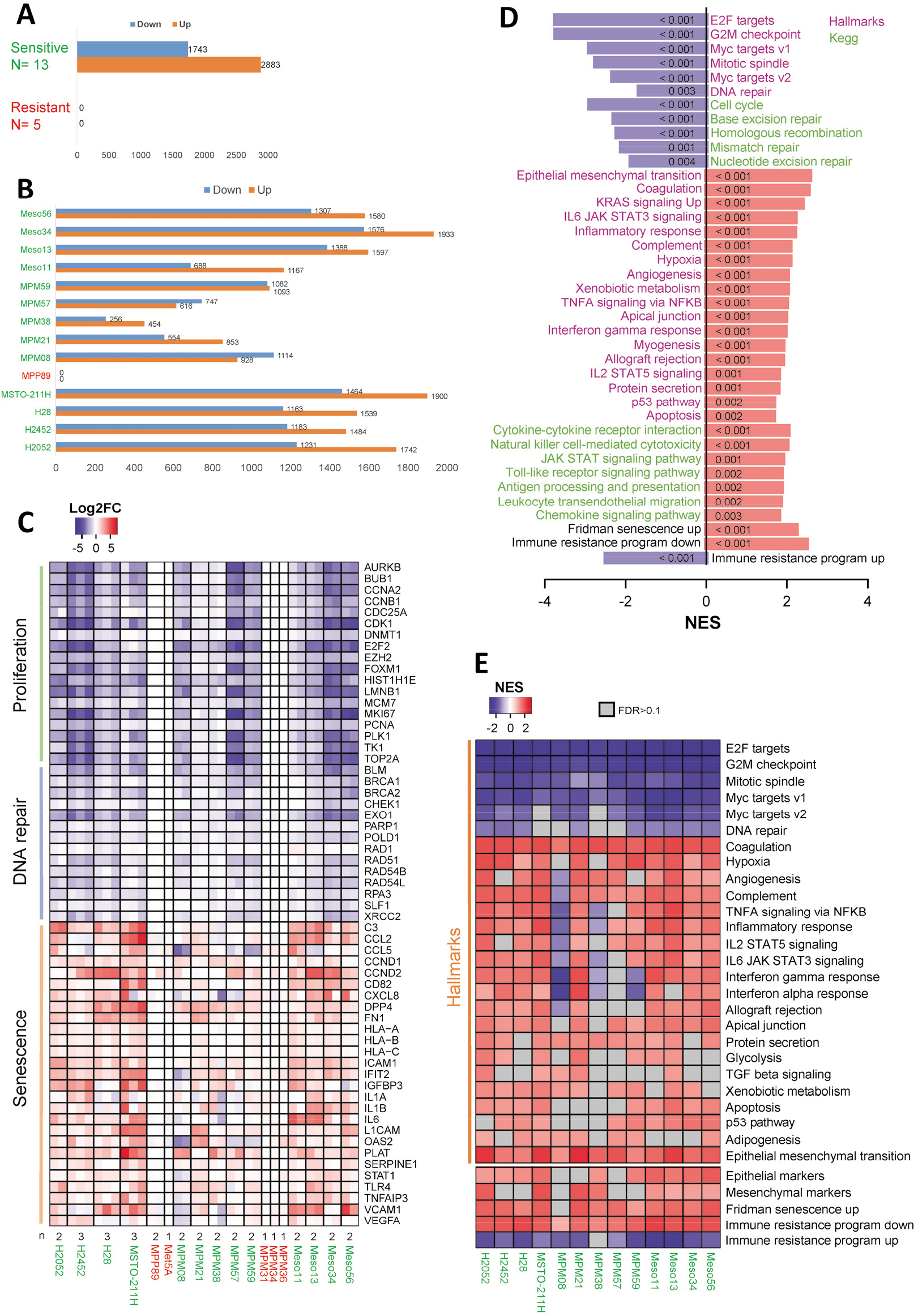
Impact of long term treatment with palbociclib analyzed by RNA-seq. 13 sensitive and 5 resistant cell lines were treated for 9-10 days with 1 μM palbociclib or DMSO. The number of independent experiments for each cell line is indicated in **C**. Sensitive cell lines are in green, resistant ones are in red. **A** Number of genes down- or up-regulated by palbociclib in a global analysis comparing sensitive and resistant cell lines. Differential expression analysis was performed using DESeq2 with min. fold change (FC) = 1.5 and false discovery rate (FDR) < 0.05. **B** Number of genes down- or up-regulated by palbociclib for each cell line with minimum 2 independent experiments (min. FC = 1.5 and FDR < 0.05). **C** Heatmap representing genes involved in proliferation, DNA repair and senescence. Data are presented as log2FC (palbo/ctrl). n independent experiments are shown, as stated below the heatmap. **D** Gene Set Enrichment Analysis (GSEA) for palbociclib-treated versus ctrl cells. Global analysis performed using all sensitive cell lines. Plot represents the normalized enrichment scores (NES) for the top regulated “Hallmarks” pathways and related “Kegg” pathways. “Fridman senescence up” and gene sets up- or down-regulated as part of an immune resistance program are also illustrated. Down-regulated pathways are in blue, up-regulated pathways are in red. FDRs obtained for each pathway are indicated in the plot bars. **E** Heatmap illustrating the “Hallmarks” pathways down- or up-regulated by palbociclib in each sensitive cell line. Epithelial/ mesenchymal markers, “Fridman senescence up” and gene sets up- or down-regulated as part of an immune resistance program are also shown. Plot represents the NES from GSEA. Pathways with FDR > 0.1 are in grey.

### Growth arrest induced by prolonged palbociclib treatment appears mostly irreversible

In clinics, palbociclib and ribociclib are administered discontinuously with one week off treatment every three weeks. Therefore, we evaluated the effect of palbociclib washout and the reversibility of the cell cycle arrest induced by this drug. When assessed by clonogenic assay, the effect of a 10-day treatment with palbociclib was mostly irreversible. This contrasted with the reversible growth arrest observed with AZD8055 and trametinib (mTOR and MEK inhibitors, respectively) for most of the investigated cell lines (Fig. 5A). However, 48 h after palbociclib washout, proliferation genes were reinduced (Fig. 5B; Supplementary Fig. S6A), which was associated with the increase of a proliferation score calculated from RNA-seq data based on the “Cell Cycle Proliferation” (CCP) signature (63) (Fig. 5C, Supplementary Fig. S6B). The phosphorylation of pRb also increased after drug withdrawal (Supplementary Fig. S4B) and cells were able to re-enter S-phase (Fig. 5D, Supplementary Fig. S6C). Nevertheless, this cell cycle re-entry might be abortive, as cells were unable to actively proliferate in the colony-forming assay. As thoroughly investigated in a recent study (64), this could be due to the accumulation of abnormal nuclear figures (micronuclei, fragmented nuclei) observed after palbociclib treatment and even more after withdrawal of the drug (Fig. 5E). Interestingly, whereas the repression of E2F-dependent cell cycle and DNA repair genes was essentially reversible, most gene up-regulations including those associated with SASP and productions of various cytokines, were maintained or even more augmented after palbociclib washout (Fig. 5B,F,G; Supplementary Fig. S4B,D, S5 and S6A,E).

**Figure 5.**
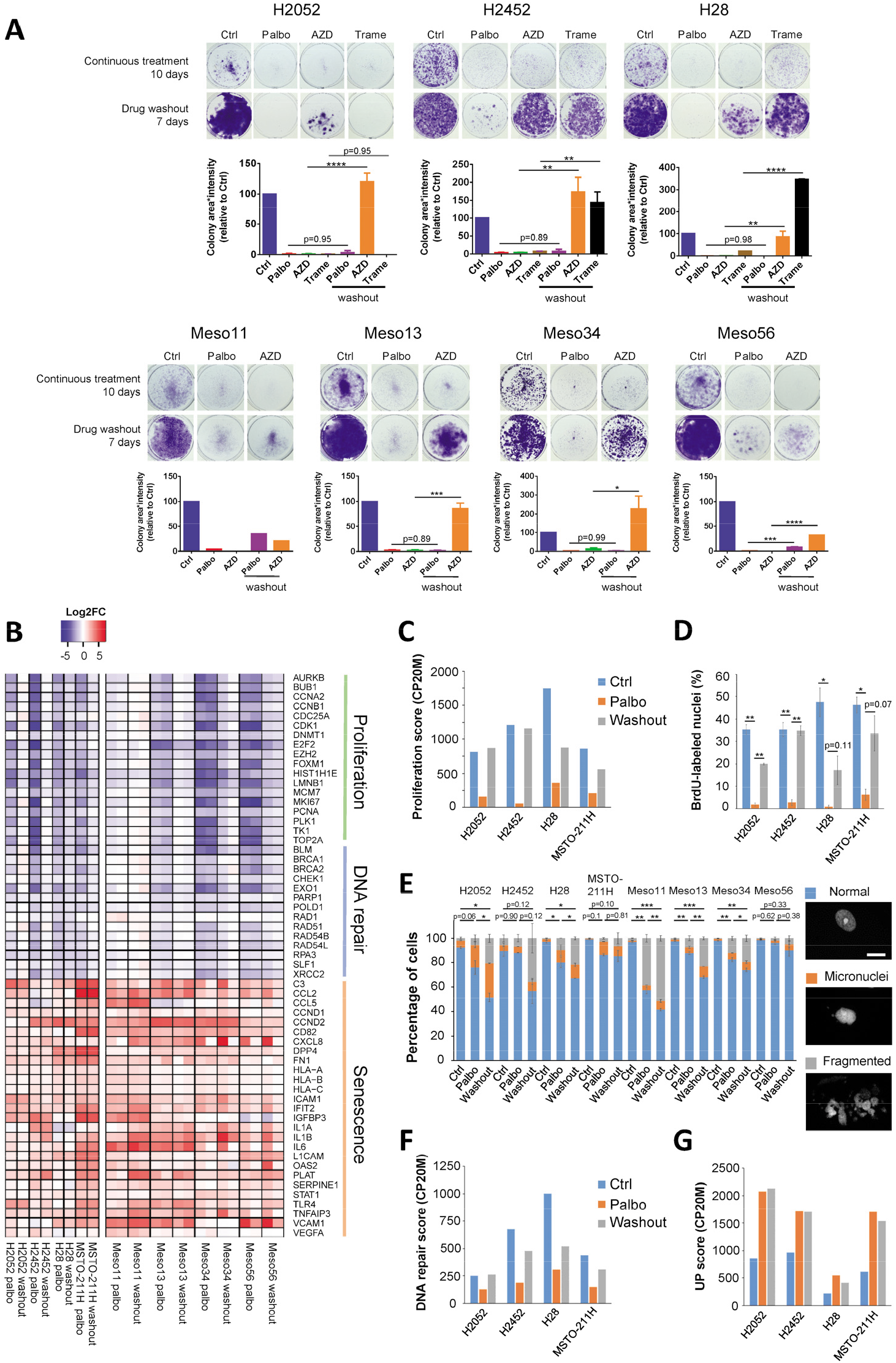
Palbociclib-induced growth arrest is poorly reversible. **A** Clonogenic assays of cells treated continuously for 10 days with 1 μM palbociclib (Palbo), 50 nM of AZD8055 (AZD, mTOR inhibitor) or 20 nM trametinib (Trame, MEK inhibitor) followed or not by a drug washout for 7 days. Representative images are illustrated above the quantification of the staining using area * intensity expressed in percentage of control (Ctrl). Mean +/− SEM of 2 independent experiments except for meso11 (n=1). **B** Differential expression of genes involved in proliferation, DNA repair and senescence assessed by RNA-seq in cells treated for 10 days with 1 μM palbociclib (Palbo) followed or not by a drug washout for 48 h. Data are presented as log2FC (Palbo/ctrl and washout/ctrl). Left part, n=1. Right part, n=2. **C** CCP proliferation score calculated from the RNA-seq data illustrated in **B** and expressed in counts per 20 million reads (CP20M). **D** DNA synthesis in cells treated as in **B**. Mean +/− SEM of 2 independent experiments. **E** Percentage of normal nuclei, micronuclei or fragmented nuclei in cells treated as in **B**. Mean +/− SEM of 2 independent experiments. Statistical analysis was performed on the abnormal nuclei (micronuclei + fragmented nuclei). Representative images are shown on the right. Scale bars, 20 μM. **F** DNA repair score calculated from the RNA-seq data and expressed in counts per 20 million reads (CP20M). This score represents the median expression of the genes involved in DNA repair illustrated in **B**. **G** Expression score of up-regulated genes calculated from the RNA-seq data illustrated in Supplementary Fig. S6A and expressed in counts per 20 million reads (CP20M). **A, D, E** *p<0.05, **p<0.01, ***p<0.001, ****p<0.0001 (ANOVA).

### CDK4 phosphorylation is detected in a majority of MPM tumors

To evaluate the proportion of MPM patients that would be potentially responsive or intrinsically resistant to CDK4/6 inhibition, we determined the CDK4 modification profile from frozen MPM tumors and normal pleura samples (Supplementary Table S7A) as investigated previously in breast cancer (48). Whereas it was absent in normal pleura, CDK4 phosphorylation was detected with variable abundance in 39 of 47 tumors, suggesting that about 80% of patients could respond to CDK4/6 inhibition (Fig. 6A; Supplementary Fig. S7). Tumors were classified as done previously (48) in three groups based on their CDK4 phosphorylation status: A (Absence of phosphorylation), L (Low phosphorylation) and H (High phosphorylation). When CDK4 phosphorylation was detected, its relative abundance significantly correlated with the “Cell Cycle Proliferation” score calculated from RNA-seq data (Fig. 6B), which was significantly different in class H versus class L tumors (Fig. 6C). Two samples with no CDK4 phosphorylation (E8, L7) were not proliferative (Fig. 6C, Supplementary Table S7A) and stained positive for p16 (Fig. 6D; Supplementary Table S7A), indicating that these tumor samples could be in a senescent state. We also note that these two patients are long-term survivors with one having a 13-year survival and the other being alive more than 6 years after diagnosis (Supplementary Table S7A). Unexpectedly, CDK4 phosphorylation was undetectable in six proliferative tumors. For four of them, this was associated with an extremely high expression of p16 at both protein (Fig. 6D; Supplementary Table S7A) and RNA (Fig. 6E; Supplementary Table S7B) levels. Interestingly, the tumor with the highest degree of CDK4 phosphorylation (E3, a patient who died two months after diagnosis) presented also an elevated *CDKN2A* mRNA expression (Fig. 6E; Supplementary Table S7B), but its p16 staining was weak (Fig. 6D; Supplementary Table S7A). This was associated with a partial *CDKN2A* deletion affecting the exon 3 (Supplementary Table S7B). High p16 expression is often associated with an inactivation of pRb/*RB1*. However, the p16-high class A tumors were not distinguished by low *RB1* expression (Supplementary Fig. S8A) and no mutation (substitution or Indel) was detected using the RNA-seq data (Supplementary Table S7B). RNA-sequencing rather revealed splicing alterations (Fig. 6F). Targeted DNA-sequencing confirmed the absence of *RB1* mutations and demonstrated the loss of one *RB1* allele in at least three of these four samples (Supplementary Fig. S8B,C). Moreover, different partial deletions of *RB1* were found in the remaining allele (Supplementary Fig. S8D,E). Our data thus suggest a double hit inactivation of the *RB1* gene in the p16-high class A tumors. Surprisingly, phospho-CDK4 was also undetectable in two proliferative tumors expressing very low levels of *CDKN2A* (E7 and L14, Supplementary Fig. S7 and Table S7B). Even though we cannot totally exclude a loss of CDK4 phosphorylation during sample processing, these two samples could represent particular class A tumors. E7 tumor has the highest expression of *CDK6* associated with high levels of *CDKN2C* (Supplementary Table S7B). This suggests it could depend on CDK6 instead of CDK4 and therefore be resistant to palbociclib, since CDK6 overexpression is a well-known mechanism of resistance to CDK4/6 inhibition (65, 66). Moreover, CDK6 up-regulation has been shown to induce *CDKN2C* encoding p18 that binds to CDK4/6, potentially preventing phosphorylation of these kinases and also competing with CDK4/6 inhibitory drugs (66). L14 tumor also expressed high levels of *CDK6* and *CDKN2C.* That, together with a low *CCND1* expression, could explain the absence of phosphorylated CDK4 in this sample. Finally, profile A tumors, including those with *RB1* deletion, were observed in the different histotypes (Fig. 6G); however *RB1* defects might be less frequent in epithelioid tumors (1 of 38 in the present cohorts).

**Figure 6.**
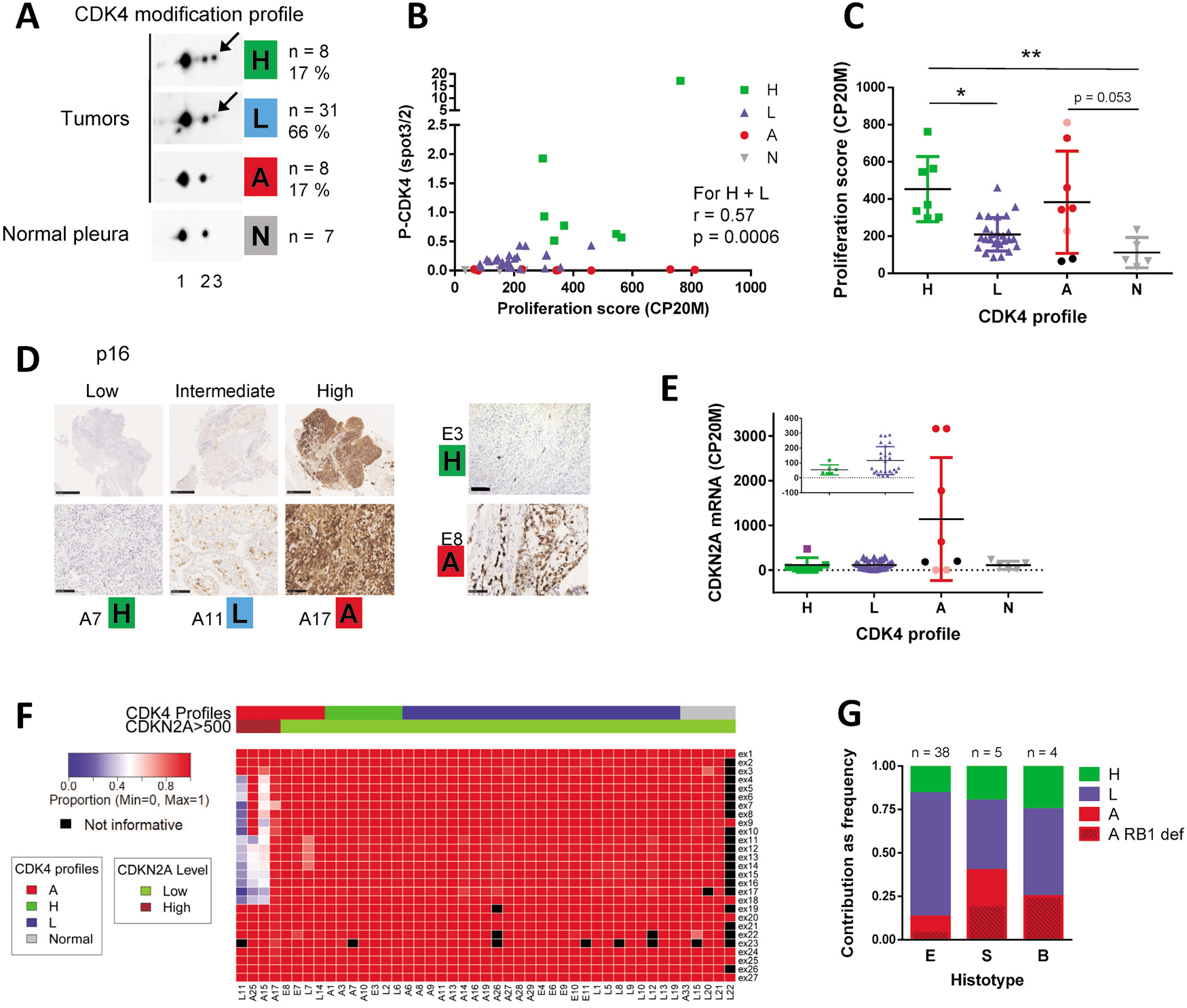
CDK4 phosphorylation is detected in a majority of MPM tumors. **A** Representative immunodetections of CDK4 separated by 2D-gel electrophoresis from 47 tumors and 7 normal pleurae. A CDK4 modification profile was attributed to each tumor based on the ratio (r) spot3/spot2 quantified from the immunoblots. High (H): r > 0.5, Low (L): 0.025 < or = r < or = 0.5, Absent (A): r < 0.025. The number and percentage of samples in each class is indicated on the right. Arrows indicate the position of the T172-phosphorylated form of CDK4 (spot 3). **B** Phosphorylation of CDK4 (ratio spot3/spot2) as a function of the CCP proliferation score calculated from RNA-seq data (n=40). Correlation between CDK4 phosphorylation and proliferation for H and L profiles was evaluated with a Spearman’s rank test. **C** Proliferation score in the different classes of tumors (H, L and A) and in normal pleura (N). For profiles A: red dots = absence of CDK4 phosphorylation with high *CDKN2A* expression and high proliferation score, pink dots = absence of CDK4 phosphorylation with low *CDKN2A* expression and high proliferation score, black dots = absence of CDK4 phosphorylation with intermediate *CDKN2A* expression and low proliferation score. Mean +/− SD, *p<0.05, **p<0.01, Kruskal-Wallis followed by Dunn’s multiple comparison tests. **D** Representative IHC staining of p16. H, L, A, CDK4 modification profile. Scale bars, 100 μM except for the three left upper panels (1 mm). **E** *CDKN2A* mRNA levels (RNA-seq data in CP20M) in the different classes of tumors and in normal tissues. For class A tumors, dots color like in **C**. In class H tumors, the purple square represents the E3 sample that presents relatively high levels of *CDKN2A* together with low p16 staining associated with a *CDKN2A* mutation. Error bars: mean +/− SD. **F** Heatmap representing the proportion of correctly spliced exons in *RB1* calculated from RNA-seq data for 40 tumors and 5 normal pleurae. Tumors are grouped by CDK4 modification profile and separated based on *CDKN2A* expression. Values were considered not informative when less than 10 reads were covering the region. **G** Relative proportions of the three CDK4 modification profiles in the different histotypes. E: epithelioid, S: sarcomatoid, B: biphasic. Class A samples with *RB1* defect are distinguished by a striped pattern. The number of samples in each histotype is indicated at the top of each bar.

## Discussion

In this work, we have evaluated the effect of CDK4/6i in a panel of 28 MPM cell lines including 19 patient-derived cell lines from two independent collections. Extending the observations by Bonelli and coll. (49) and Aliagas and coll. (50), we found that the majority of these models were particularly sensitive to CDK4/6 inhibition, irrespective of their histotype. Indeed, in most cell lines, continuous palbociclib treatment produced an almost complete cell cycle inhibition, which was sustained for at least two weeks. This prolonged treatment irreversibly inhibited the proliferation in colony-forming assay, despite re-induction of pRb phosphorylation and cell cycle genes upon drug washout. A SASP including various potentially immunogenic components was also irreversibly induced. This remarkable sensitivity of MPM cell lines to CDK4/6 inhibition contrasts with a more partial response and/or rapidly developing resistance (adaptation) observed in other cancers such as estrogen receptor-positive breast cancer (58), pancreatic ductal adenocarcinoma (PDAC) (67, 68), colorectal cancer (69) and anaplastic thyroid cancer (70). In those tumors, combined inhibition of CDK4/6 and signaling pathways upstream of CDK4 activation (MAPK and/or PI3K-mTOR (43)) is required to fully and durably suppress their proliferation (58, 67, 68, 71, 72). Such an exquisite sensitivity to CDK4/6 inhibition might be explained by the absence of oncogenic driver mutations affecting those signaling cascades in mesotheliomas, which are mostly characterized by the inactivation of tumor suppressor genes (6, 10–12).

On the other hand, four MPM cell lines and immortalized MeT-5A cells were completely insensitive to CDK4/6i, with no inhibition of cell cycle progression and no induction of SASP transcriptional responses. For most of these cell lines, the insensitivity to palbociclib was associated with their lack of T172-phosphorylation of CDK4 and high p16 accumulation that diverted CDK4 from binding D-type cyclins. However, *RB1* defects could neither be detected by RNA-seq nor by analyses of pRb accumulation and phosphorylation in the patient-derived cell lines MPM_31 and MPM_34. Therefore, superior to pRb, the status of CDK4 phosphorylation was able to predict the response to palbociclib in 27/28 mesothelial cell lines, as we also observed it in breast cancer (48). The involvement of p16 is further supported by the observation that the only resistant cell line displaying some CDK4 phosphorylation (MPM_36) combines a deletion of *RB1* with the loss of *CDKN2A.* Among other causes (73), p16 over-expression in cancer cells has been related to *RB1* loss of function (52, 53), which prevents the *CDKN2A* locus silencing by Polycomb Repressor Complexes (74, 75) and trimethylation of lysine 27 on histone H3 by EZH2 (76, 77). This silencing might be directed by interaction of EZH2 with pRb and E2F1 (78). Interestingly, our data suggest that high levels of *p16/CDKN2A* could also be associated with constitutive over-activation of CDK2 in MPM_31 and MPM_34 cells, resulting from amplification of *CCNE1* and possibly associated with other mechanisms including a *p27/CDKN1B* mutation or high cdc25 phosphatase activity (Fig. 2E-G). This over-activation of CDK2 can result in constitutive inactivation of pRb by phosphorylation. Alternatively, we do not exclude that these high p16 levels could be the consequence of replicative stress (79), which might be provoked by deregulated CDK2 activity. Indeed, both *CCNE1* over-expression and *RB1* alterations have been shown to induce the DNA Damage Response (DDR) (80). Our results thus suggest that in cells with wild-type *CDKN2A,* the insensitivity to CDK4 inhibition could be directly due to the absence of the main target of CDK4/6i, i.e. active phosphorylated CDK4, rather than to the two generally considered mechanisms - absence/mutation of pRb, or amplified cyclin E1-mediated activation of CDK2 (33, 58, 81–83). It remains difficult to understand why the deficiency of pRb is sufficient to generate the observed insensitivity to CDK4/6i in the presence of the other proteins of the RB family, p107 (*RBL1*) and p130 (*RB2*), which are also inactivated by CDK4 phosphorylation and capable of blocking the cell cycle (34, 35, 37, 38, 84, 85). The absence of CDK4 phosphorylation is a more likely explanation for a complete insensitivity to CDK4 inhibitors. Indeed, as the T172-phosphorylation is required for the opening of the catalytic site of CDK4 (86, 87), it is not only critical for the activity of CDK4 complexes, but also potentially required for the engagement of CDK4 by ATP-competitive inhibitors like palbociclib. This likely explains why in RB-deficient models expressing high p16, palbociclib is observed not to bind to CDK4 (88, 89). The lack of activating T177-phosphorylation in CDK6, which is more frequent (44), may also explain the weak binding of CDK4/6i to CDK6 in the resistant cells that over-express it (66, 90). Indeed, the sensitivity to CDK4/6i was restored by the S178P mutation of CDK6 (90) that forces its T177-phosphorylation (44).

We detected CDK4 phosphorylation in 80 % of MPM tumors suggesting that the majority of the patients could at least initially respond to CDK4/6i. Treatment of relapsed mesothelioma with abemaciclib has resulted in 15 % partial responses and 54 % stable diseases in a recent phase II clinical trial [NCT03654833] using negative p16 IHC staining as an inclusion criterion (51). However, as initially observed in a subset of breast cancers (48) and here in most insensitive MPM cell lines, 15 % of MPM tumors were highly proliferative despite the lack of CDK4 phosphorylation and thus are expected to be intrinsically resistant to CDK4/6i. For two thirds of them, this was associated with copy number loss and/or partial deletions of *RB1* leading to a high p16 expression, which has also been observed by others in 17% of 88 MPMs (91). Whether such elevated p16 accumulations might be associated to previously unrecognized *RB1* alterations including some monoallelic deletions, which were reported to be relatively frequent (up to 25 %) (15, 92), or other causes such as inactivation of pRb by CDK2-dependent phosphorylation, should be evaluated in larger MPM cohorts. If frozen tissue samples are available, the presence or absence of phosphorylated CDK4 could be the most direct biomarker to predict sensitivity to CDK4/6i. However, in FFPE samples, the IHC detection of phosphorylated CDK4 would be complicated by its extremely low abundance (93) and by the loss of phosphorylation events before and during formalin fixation. Gene expression signatures including from RNA-seq data might instead be used to predict the CDK4 status (48). Since most MPM tumors lacking phospho-CDK4 had high p16 expression, appropriately scored p16 immunohistochemistry (especially regarding intensity and homogeneity of expression (53, 91)), possibly in combination with proliferation markers, could also be used to exclude those intrinsically resistant tumors.

*RB1* status has been shown to govern differential sensitivity to genotoxic and molecularly targeted therapeutic agents, *RB1* defects generally sensitizing to the formers (94, 95). Interestingly, in tumors predicted to be insensitive to CDK4/6i, transient administration of these drugs could be used to transiently arrest proliferation of normal cells and prevent chemotherapy-induced myelosuppression, hence allowing to increase the dose of genotoxic drugs such as cisplatin and thus the therapeutic window between normal and transformed cells (96). This idea has been validated clinically in small cell lung cancers that are generally insensitive to CDK4/6i due to their *RB1* locus alteration (97), leading to recent approval by the FDA of trilaciclib for this indication. Therefore, defining or predicting the CDK4 phosphorylation status might really be one key to tailor the use of CDK4/6i in MPM treatment and their combination with targeted or genotoxic therapies.

As characterized in some other cancers (21, 25, 31, 98), our present data suggest that MPM might be especially responsive to various (non-exclusive) combinatorial approaches to extend the cytostatic activity of CDK4/6i into a real clearance and eradication of tumor cells:

1. *Combination with senolytics.* Senescent cells harbor vulnerabilities allowing their specific killing by various ‘senolytics’ (23, 99), including BCL-2 and BCL-xL inhibitors (venetoclax, navitoclax, A1331852,...) that were observed to augment the tumor response to CDK4/6i (100, 101). Prolonged palbociclib treatment indeed increased transcriptomic pathways related to apoptosis in 9 of 13 sensitive MPM cell lines and apoptotic cell death induced by the BCLi Navitoclax and A1155463 (Fig. 3C). Further studies should evaluate new generations of such drugs, e.g. recently described BCL-XL PROTAC degraders (102) or galacto-conjugated navitoclax prodrug to specifically target the SAβGal-positive senescent cells (103).
2. *Combinations of CDK4/6i with chemotherapy.* CDK4/6 inhibition, by blocking cells into G1 phase, is generally thought to antagonize the effect of chemotherapy. However, several clues and recent reports suggest a high efficacy of sequential combinations (25, 26): (i) in clinics, palbociclib and ribociclib are used discontinuously with one week off treatment every 3 weeks. We observed that palbociclib paradoxically stabilizes p21/p27-free cyclin D-CDK4/6 complexes that become hyperactive upon palbociclib withdrawal, potentially inducing a burst of tumor cell cycle progression that might open a window of increased responsiveness to genotoxic therapies (104). As thoroughly investigated in a recent study (64), we observed here that following a prolonged G1-arrest, the cell cycle entry induced by palbociclib washout is clearly perturbed, as it did not lead to clonogenic proliferation but to various potentially stressful nuclear defects; (ii) we observed that palbociclib down-regulates various DNA repair genes and pathways in MPM cells. As reported in other models, this might prevent DNA repair by homologous recombination and cell recovery after chemotherapy, which would favor a cooperation between CDK4/6i and DNA repair inhibitors such as PARP inhibitors (olaparib) (64, 105–107). Moreover, therapy-induced SASP increases delivery of chemotherapy by inducing a vascular remodeling (108), which might also occur in CDK4/6i-treated MPM, as suggested in our data by the up-regulation of angiogenesis pathway and VEGF in 9 of 13 sensitive MPM cell lines.
3. *Combination of CDK4/6i with immunotherapy.* Inhibition of CDK4/6 has been shown to favor the elimination of tumor cells by the adaptive immune system through various mechanisms including enhanced antigen presentation by cancer cells, increased infiltration and activation of T cells; and inhibition of regulatory T cells proliferation (27, 28, 109–111). On the other hand, therapy-induced senescence was also shown to elicit a distinct mechanism of innate immune attack by NK cells, which is mediated by an NF-kB–dependent SASP program that culminates in the secretion of pro-inflammatory cytokines and surface expression of NK cell-activating molecules such as ICAM-1 (67). In our RNA-seq data, pathways related to immune response and antigen presentation were among the top up-regulated ones in response to palbociclib. CDK4/6 inhibition was associated with induction of a SASP of variable composition in the different cell lines. This included ICAM-1 and pro-inflammatory cytokines/chemokines such as IL6/CCL2/CCL5, known to promote the recruitment of various immune cells (24, 67, 112). Moreover, we observed that CDK4/6 inhibition represses a cancer cell program shown to mediate resistance to anti-PD1 therapy in melanoma, consistent with studies demonstrating a cooperation between CDK4/6i and immune checkpoint blockade (62, 113). Interestingly, after palbociclib washout most gene up-regulations associated with SASP including productions of various cytokines were maintained or even more augmented (at variance with cell cycle genes), as also recently observed by others (112). Therefore, this raises the plausible expectation that the drug-holiday periods of patient treatments with palbociclib and ribociclib, in complement of inducing important stressful nuclear perturbations associated to unbalanced reinduction of cell cycle genes, would sustain or enhance immune responses against tumor cells generated by the various SASP components. Overall, CDK4/6i, in addition to their direct cytostatic tumor action, would greatly facilitate response of MPMs to immune checkpoint inhibitors including the newly FDA-approved combination of nivolumab and ipilimumab (4, 9, 113).

To conclude, our study supports further clinical evaluation of CDK4/6i for the treatment of pleural mesotheliomas including in various combinations with the standard therapies. Most MPM patients could respond to CDK4/6i, which may not only arrest tumor growth but also help to convert the mesotheliomas that are rarely immunologically ‘cold’ but more frequently in intermediate inflammatory states (114), into ‘hot’ tumors responsive to immunotherapy. Nevertheless, a minority of intrinsically CDK4/6i insensitive tumors lacking CDK4 phosphorylation might have to be identified.

## Materials and methods

The antibodies and drugs used in this work are listed in Supplementary Table S8.

### Cell culture

MeT-5A, NCI-H2052, NCI-H2452, NCI-H28 and MSTO-211H were obtained from ATCC (Manassas, VA, USA), MPP89 from Interlab Cell Line Collection (Genoa, Italy), DM-3 and RS-5 from DSMZ (Brunswick, Germany), and NCI-H2369 from Wellcome Sanger Institute. These authenticated cell lines were passed for fewer than 4 months after receipt. Meso11, Meso13, Meso34 and Meso56 were established in the laboratory of Marc Gregoire and Christophe Blanquart (CRCI2NA, Nantes) from pleural effusions from patients who had not received any chemotherapy (115). They were characterized phenotypically for genetic alterations in key genes of mesothelial carcinogenesis including *CDKN2A, CDKN2B, BAP1, NF2, LATS2* and *TP53* using a targeted sequencing described in (116) and karyotyping (GSE134349) (60). MPM_04 to MPM_66 cell lines are primary mesothelioma cell lines obtained from surgical resections, pleural biopsies or malignant pleural fluids of confirmed MPM cases in the research teams of Marie-Claude Jaurand and Didier Jean (Centre de Recherche des Cordeliers, Paris). They were previously characterized for genetic alterations using the same targeted sequencing (116) and some of them were characterized for copy number variations by Singlenucleotide polymorphism (SNP) arrays. The patient-derived cell lines were used in several studies showing their relevance to MPM primary tumors (7, 61, 115, 117, 118). These cell lines were authenticated based on specific gene mutations. DM-3 and RS-5 cells were cultured in NCTC-109 medium (Gibco, Carlsbad, CA, USA) supplemented with antibiotics, sodium pyruvate (1mM), glutamine (2mM), N-Acetyl-L-cysteine (1mM, Sigma Aldrich, St Louis, MO, USA) and 10% (for RS-5) or 20% (for DM-3) FBS (Gibco). All the other cells were cultured in RPMI medium (Gibco) supplemented with antibiotics, sodium pyruvate (1mM) and 10% FBS.

### Human MPM samples

Fresh frozen mesothelioma and normal pleura samples were obtained from Biobanque of Hôpital Erasme (ULB), French MESOBANK (Centre Léon Bérard, Lyon), or UZA Tumor bank, Antwerp University Hospital, Belgium. For a prospective study, MPMs were resected at the Thoracic Surgery Department of Hôpital Erasme (n=4). This study was approved by the ethics committees of Jules Bordet Institute, Erasme academic hospital and UZ Antwerpen. Tissue samples from French MESOBANK were collected in agreement with all applicable laws, rules, and requests of French and European government authorities, including the patient’s informed consents. These samples were prepared by BB-0033-00050, CRB Centre Léon Bérard, Lyon France.

### DNA synthesis

DNA-replicating cells were identified from duplicated dishes by incubation with bromodeoxyuridine (BrdU) for 1 h and counted at the microscope as described (119). For the slow-growing RS-5 cells, BrdU incubation was done for the last 24 h of a 48 h treatment with palbociclib. Alternatively, cells were seeded in triplicates in 96-well plates and BrdU was replaced by EdU. Plates were processed as described in Supplementary methods.

Pictures of abnormal nuclei were acquired using an epifluorescence microscope Zeiss Axioplan 2 (Carl Zeiss, Oberkochen, Germany) equipped with a Spot RT3 camera and the Spot 5.2 imaging software (Spot imaging, Sterling Heights, MI, USA).

### Cell growth assays

Cells were seeded in triplicates in 96-well plates and treated the day after with serial dilutions of CDK4/6i (palbociclib, ribociclib or abemaciclib) for 48 h (MTT assay) or 144 h (Sulforhodamine B (SRB) assay). MTT and SRB assays were performed as described (48).

### Clonogenic assays

5 x 10^3^ cells were seeded in 6-well plates and treated the day after with the indicated drugs for 10 days. For drug washout, cells were washed twice with PBS before incubation in complete medium for 7 days. After removal of culture medium, cells were washed with PBS and fixed in a 10% formalin solution (Sigma-Aldrich, St. Louis, MO, USA) for 10 min. Cells were washed again with PBS and stained for 30 min with 0.05 % crystal violet solution (Sigma-Aldrich), washed thoroughly, and airdried. Photographed pictures of the plates were quantified with the “ColonyArea” plugin (120) in ImageJ software.

### SA-β-gal activity assay

5 x 10^3^ cells (or 10^3^ for the control) were seeded in 6-well plates and treated the day after with DMSO or 1 μM palbociclib for 9 days. SA-β-gal activity was measured using the SA-β-gal staining kit (Cell Signaling Technology, Danvers, MA, USA) according to the manufacturer’s protocol. After overnight incubation at 37°C without CO2, the percentage of SA-β-gal positive cells was determined by counting at least 500 cells per well.

### Cytokine array

3 x 10^5^ to 10^6^ cells were plated in 9 cm Petri-dishes and treated with DMSO or 1 μM palbociclib for 9-10 days. Cells were trypsinized when necessary and medium was changed every 3-4 days. Culture supernatants were collected 48 or 72 h after the last medium replacement and cells were lysed to perform immunoblotting and allow normalization of media based on protein quantification. Supernatants were sent to Tebu-Bio (Le Perray en Yvelines, France) for analysis on a custom Quantibody Multiplex Elisa Array (RayBiotech, GA, USA).

### Immunoprecipitations and pRb-kinase assay

Co-immunoprecipitations were performed as described (42, 121). pRb-kinase activity of immunoprecipitated CDK complexes was measured by in vitro incubation with ATP and a fragment of pRb, as described (104, 121).

### Western blots

Equal amounts of whole cell extract proteins or immunoprecipitates were separated by SDS-PAGE and immunodetected.

For 2D-gel electrophoresis, cells were lysed in a buffer containing 7 M urea and 2 M thiourea. Frozen tumor slides (minimum 7 sections of 7-μm per sample) or frozen tissue powder obtained by cryogrinding were solubilized as described (48). Proteins were separated by isoelectric focusing on immobilized linear pH gradient strips pH 5-8 for CDK4 (BioRad, Hercules, CA, USA) or pH 3–10 for CDK2 (Amersham Biosciences, GE Healthcare Europe, Diegem, Belgium) before separation by SDS-PAGE (42, 57).

Chemiluminescence detections were captured on films or with a Vilber-Lourmat Solo7S camera and quantified using the Bio1D software (Vilber-Lourmat, Marne-la-Vallée, France). The profile of CDK4 separated by 2D-gel electrophoresis has been characterized previously (42, 48). The most basic form (spot 1) corresponds to unmodified CDK4. The most acidic form (spot 3) had been identified as T172-phosphorylated CDK4 form using several approaches including a T172-phosphospecific antibody. Another yet unidentified modified CDK4 form (spot 2) does not incorporate [32P] phosphate. The background-subtracted volume ratio (spot 3/spot 2) was used to define the type of CDK4 modification profile of the tumors (48). A profile A (Absent) was attributed to the sample when its ratio was below 0.025. A profile L (Low) was attributed to the sample if this ratio was between 0.025 and 0.5, while a profile H (High) was given for ratio above 0.5.

### Immunohistochemistry

Immunohistochemical stainings of Ki-67 and p16 were performed using standard routine protocols and scored by a pathologist. Pictures were acquired with a Moticam Pro camera connected to Motic AE31 microscope or a NanoZoomer digital scanner (Hamamatsu Photonics, Hamamatsu, Japan) at 40x resolution.

### RNA-sequencing

Total RNA was isolated from cell lines or frozen tumor tissues using the RNeasy Mini Kit according to the manufacturer’s protocol (Qiagen, Hilden, Germany). RNA yield and purity were assessed using a Fragment Analyzer 5200 (Agilent technologies, Massy, France). 100 ng of RNA was used to create indexed cDNA libraries using the NEBNext Ultra II Directional RNA Library Prep Kit for Illumina (New Englands Biolabs, Ipswich, MA, USA) following manufacturer’s protocol. The multiplexed libraries were loaded on a NovaSeq 6000 (Illumina, San Diego, CA, USA) using a S2 flow cell and sequences were produced using a 200 Cycles Kit. Paired-end reads were mapped against the human reference genome GRCh38 using STAR software to generate read alignments for each sample. Annotations Homo_sapiens.GRCh38.90.gtf were obtained from ftp.Ensembl.org. After transcripts assembling, gene level counts were calculated with HTSeq and normalized to library size to obtain counts per 20 million reads (CP20M).

### Analysis of RNA-seq data

The CP20M count matrix was filtered to keep genes expressing a minimum of 100 counts in at least one sample. Differentially expressed genes with a fold change (FC) > 1.5 or < 1/1.5 and false discovery rate (FDR) < 0.05 were identified using DESeq2 in R or in the iDEP.91 platform. The model ‘treatment+paired’ was used to allow pairwise comparison between the treated and control samples within each experiment. For pathway analysis, the ranked gene list was used in Gene Set Enrichment Analysis (GSEA) Preranked using the “Hallmarks” and “Kegg” Molecular Signatures database (MSigDB) gene sets v7.2. Fridman_senescence_up (from the “Curated” gene sets), gene sets containing epithelial and mesenchymal markers previously identified in MPM (61) and the gene sets up- and down-regulated as part of the immune resistance program described by Jerby-Arnon and coll. (62) were also used. Heatmaps were generated using the heatmap.2 package in R. A proliferation score was calculated by using the median expression of the genes of the Cell Cycle Progression (CCP) signature described in (63).

### Analysis of splicing junctions

The proportions of appropriately spliced exons were estimated using a custom script working in R, as detailed in supplementary methods. A heatmap of these proportions was drawn with the heatmap.3 function from the heatmap3 package. If the number of reads for a given exon is below 10, the data are considered as not informative and appear as a black cell in the heatmap.

**Single-nucleotide polymorphism (SNP) arrays and qRT-PCR** are detailed in Supplementary methods.

### Targeted DNA-sequencing

Genomic DNA was extracted from frozen tissue using the QiaAmp Mini Kit or the DNA/RNA extraction mini kit from Qiagen. DNA was quantified and quality checked with the Qubit fluorometer or the Quant-iT PicoGreen dsDNA Assay Kit (Thermo Fisher Scientific, Waltham, MA, USA). Massive parallel sequencing was performed using targeted-capturing of the 165 genes included in the “Solid and Haematological tumors” panel (BRIGHTCore, Brussels, Belgium). 150 ng of genomic DNA was fragmented and processed to construct libraries with barcodes, which were hybridized with the DNA panel. The libraries were sequenced on Illumina NovaSeq 6000 with a coverage of 1500 x. Copy number variation was analyzed as detailed in Supplementary methods.

### Analysis of mutations

Integrative Genomics Viewer software (IGV version 2.7.2, Broad Institute) was used to visualize mutations from RNA-seq and targeted DNA-sequencing.

### Statistical analysis

Statistical analysis was performed using GraphPad Prism 6 (GraphPad Software, La Jolla, CA, USA). The two-sided unpaired Student’s t test was used for comparison between two groups. Multiple group comparisons were done using one-way ANOVA followed by Holm-Sidak’s multiple comparison tests or the Kruskal-Wallis test followed by Dunn’s multiple comparison tests, as stated in the figure legends. Correlation was evaluated with a Spearman’s rank test.

## Data availability

The RNA-sequencing data from commercial cell lines and the SNP array data from patient-derived primary cell lines have been deposited in the Gene Expression Omnibus with accession number: GSE195568 and GSE197288, respectively. The RNA-sequencing data from patient-derived cell lines and the data from MPM patients will be deposited at the European Genome-phenome Archive (EGA), which is hosted by the EBI and the CRG, under accession number EGAS00001006117.

## Funding details and acknowledgements

This study was supported by the Belgian Foundation against Cancer (grants 2014-130 and 2018-138); the Fonds de la Recherche Scientifique-FNRS (FRS-FNRS) under Grants J.0002.16, J.0141.19 and J.0169.22); Télévie (grant 7.4589.17); WALInnov 2017.2 (CICLIBTEST 1710166); and the Fund Doctor J.P. Naets managed by the King Baudouin Foundation. XB received fundings from the European Union’s Horizon 2020 research and innovation program under the Marie Skłodowska-Curie grant agreement No 843107 and from the Région de Bruxelles Capitale – Innoviris (RBC/BFB1). MT and PPR are (respectively) Posdoctoral Researcher and Senior Research Associate of the FRS-FNRS.

The work performed at Centre de Recherche des Cordeliers was supported by Inserm, the Ligue Contre le Cancer (Ile de France committee and national program Cartes d’Identité des Tumeurs (CIT)), the Chancellerie des Universités de Paris (Legs POIX) and grants from Fondation pour la Recherche Médicale (FRM, to JBA) and from Cancéropôle Ile-de-France. The work performed at CRCI2NA was supported by Inserm, CNRS, the Ligue Contre le Cancer (committees of Morbihan, Sarthe, Vendée et Loire-Atlantique) and ARSMESO44.

We thank the Biobanque of Erasme Hospital (ULB) (especially Flavienne Sandras) and UZA Tumor bank (Antwerp University Hospital), which are both funded by the Belgian National Cancer Plan. MESOPATH and MESOBANK data and samples collection are supported by the French National Cancer Institute (INCA) core grant and the National Health Institute (Santé Publique France). We also thank the Brussels Interuniversity Genomics High Throughput Core (www.brightcore.be) and Dr Anne Lefort for RNA handling and sequencing.

We thank Professors Jacques Dumont, Marie-Claude Jaurand and Marc Grégoire for continued interest and support; Prof. Paul de Vuyst and Georgina Amand for initial participation and communication of clinical data; Marieke Hylebos and Lisa Quetel for participation in initial experiments; Daniel Pouliquen for initial discussions and useful information related to meso cell lines; Jaime M. Pita and Katia Coulonval for discussion and critical reading of the manuscript; and Vincent Vercruysse for technical assistance.

## Contributions

PPR and ER conceived the project; SP, CM and JBA designed and performed experiments; PPR, ER, SP, XB, CM, CB and DJ analyzed and discussed the data; ER, MT, YB and FL performed bioinformatics analyses; CB and DJ provided patient-derived cell lines; MR, PP, NLS, STE, FGS, JPVM and TB provided MPM samples and related data; SP and PPR wrote the original draft. All the authors contributed to manuscript editing and approved the final version.

