## Supplemental methods and figures for "Preclinical evaluation of CDK4 phosphorylation predicts high sensitivity of malignant pleural mesotheliomas to CDK4/6 inhibition"

10 Present address: IGDR UMR 6290, CNRS, Université de Rennes 1, Rennes, France.

#### This pdf file contains the supplementary methods, references and figures (S1-S8).

**Supplementary tables are in excel files:**

**One combined excel file for:**

- **Table S1.** Cell lines characteristics
- **Table S2.** Quantification of viral genomes from RNA-seq data
- **Table S3.** Genes down- or up-regulated after long term treatment with palbociclib (global analysis)
- **Table S5.** Differentially expressed pathways after long term treatment with palbociclib in a global GSEA analysis: **A** Down-regulated pathways, **B** Up-regulated pathways
- **Table S7.** Tumors characteristics: **A** Clinical data, **B** RNA-seq data
- **Table S8.** Drugs and antibodies

**One separate excel file for:**

- **Table S4.** Genes down- or up-regulated after long term treatment with palbociclib (analysis by cell line)
- **Table S6.** Pathways deregulated after long term treatment with palbociclib in each cell line

#### Supplementary methods

##### EdU Cell Proliferation Assay

Cells were seeded in a 96-well plate ($10^4$ cells/well) and, once they reached 50% confluency, treated with DMSO (control) or Palbociclib at increasing concentrations for 24 h. EdU (5-ethynyl-2'-desoxyuridine, Thermo Fischer Scientific, Waltham, MA, USA) was incorporated (10 µM final concentration) 1 h before fixation and nuclei staining using 4% formalin (Sigma-Aldrich, St. Louis, MO, USA) and 5µg/ml Hoechst 33342 (Thermo Fischer Scientific) solution for 15 min at room temperature. Permeabilization was achieved with 0.3% Triton X-100 (Sigma-Aldrich) solution for 20 min at room temperature. Cells were rinsed with 1% Bovine Albumin Fraction V (Thermo Fischer Scientific) solution. Staining was done with 1 mM Copper (II) sulfate solution (Sigma-Aldrich), 100mM L-Ascorbic acid (Sigma-Aldrich) and 3 µM Alexa Fluor™ 647 Azide (Thermo Fischer Scientific) for 30 min at room temperature. Cells were rinsed with 1% Bovine Albumin Fraction V (Thermo Fischer Scientific) solution. Nuclei counting and Edu positive cells were assessed with an Operetta High Content Imaging System (PerkinElmer, Waltham, MA, USA). All conditions were done in triplicates from two independent experiments.

##### Viral genome detection and quantification

The presence of viral genome in malignant pleural mesothelioma tumors (MPM) and cell lines was explored by aligning the raw reads using the STAR algorithm to a custom GTF file built with the genomic sequences of viruses. This GTF file was built by downloading the fasta files sequences of the viral genomes from the NCBI database and converting them into a GTF file using a dedicated script in R (available upon request). Each viral genome is recorded as a single gene in this GTF file. Gene level counts were normalized to library size to obtain counts per 20 million reads (CP20M). A threshold of minimum 20 counts was applied to minimize the rate of false-positive hits in virus detection.

##### Analysis of splicing junctions

The genomic coordinates of the *RB1* gene exons were first extracted from Ensembl with a R script extracting data corresponding to the ENSG00000139687 id using the library biomaRt. The coordinates from the introns are computed from the coordinates of the exons. The genomic position at -200 nucleotide of the transcription start site is used as the starting coordinate of the upstream sequence. The length of the exons was computed based on the coordinates of the exons. Next, for a given exon, we identify the most distant exons in 5' for which the sequence could be included after splicing in a read of 97 nucleotides ending in the given exon. The *RB1* gene exon genomic coordinates are converted to a genomic Range object using the makeGRangesFromDataFrame function of the GenomicRanges library. BAM files are opened using the BamFile function of the Rsamtools library. The function scanBam of the GenomicRanges library is used to extract the reads, their position and the cigars information. For each read falling in a genomic coordinate range corresponding to an exon for which at least 10 reads were recorded, the script identifies whether the coordinate of the start of the read is located within the expected exons located upstream of the considered exon defined above. The number of reads for which the start coordinates are within an expected exon defined for each exon and each sample is divided by the number of informative reads to compute the expected splicing ratio. Commented scripts to reproduce the data are available upon request.

**DNA-sequencing analyses**

Copy number calling. We derived genome-wide copy number profiles using the off-target reads from our targeted panel, as these provide a uniform shallow-coverage whole-genome sequencing experiment. We first remove the on-target regions from the bed definition of the target with padding of 1000bp upstream and downstream. Then we bin the rest of the genome in 30kb bins and count the number of reads with MAPQ>30 falling in each bin to obtain a read-count track genome-wide. We divide the number of reads in each bin by the average number to get a ratio r, which can be expressed as a function of the purity $\rho$, the average number of copies (or average ploidy $\psi$) and the local number of DNA copies nT: $r=(n_T \rho+2(1-\rho))/\psi$ . We take the log of this track and segment it with circular binary segmentation (1) from the R package DNAcopy v1.64.0 with default parameters. Then for each value of the ploidy $\psi \in [1.5,5]$ by 0.01 and purity $\rho \in [.2,1]$ by 0.01 we fit $n_T$ of each segment from the first equation $n_T=(\psi r-2(1-\rho))/\rho$. We calculate the sum of euclidean distances between $n_T$ and the closest integer values, i.e. round($n_T$), and then select the combination of values of $\psi$ and $\rho$ that minimizes this distance. For copy number of exons from on-target reads, we counted the number of aligned reads with MAPQ>30 at the center of each exon target and divided the counts by the counts from the identified diploid sample L2 to normalize for different target-capture efficiency.

RNA splice junctions. From the gene annotation from Ensembl v104.38, we looked at the mate coordinates of reads aligning at the exon junctions, showing abnormal amounts of non-canonical junctions in *RB1* of four of the samples.

**Single-nucleotide polymorphism (SNP) arrays**

Some MPM cell lines were characterized for copy number variations using Illumina HumanOmniExpress-24 v1.0 BeadChip SNP arrays. Integragen SA (Evry, France) carried out hybridization, according to the manufacturer's recommendations. The BeadStudio software (Illumina) was used to normalize raw fluorescent signals and to obtain log R ratio (LRR) and B allele frequency (BAF) values. Asymmetry in BAF signals due to bias between the two dyes used in Illumina assays was corrected using the tQN normalization procedure (2). We used the circular binary segmentation algorithm (3) to segment genomic profiles and assign corresponding smoothed values of log R ratio and B allele frequency. The Genome Alteration Print (GAP) method was used to determine the ploidy of each sample, the level of contamination with normal cells and the allele-specific copy number of each segment (4).

**Quantitative Real Time -PCR (qRT-PCR) analysis**

Total RNA (1.5 µg) was reverse transcribed in a final volume of 50 µl using the High Capacity cDNA Reverse Transcription kit (Thermo Fisher Scientific). qRT-PCR reactions were performed using TaqMan probes (Thermofisher) and the high throughput BioMark HD system (Fluidigm) following manufacturer's instructions. Expression data (Ct values) were acquired using the Fluidigm Real Time PCR Analysis software. The mean of the following 5 housekeeping genes *ACTB* (Hs01060665_g1), *CLTC* (Hs00964504_m1), *GAPDH* (Hs02758991_g1), *TBP* (Hs00427620_m1), *RNA18S* (Hs03928990_g1) was used for the normalization of *CDKN2A* (Hs00923894_m1) expression data (-ΔCt).

### Supplementary Figure S1A

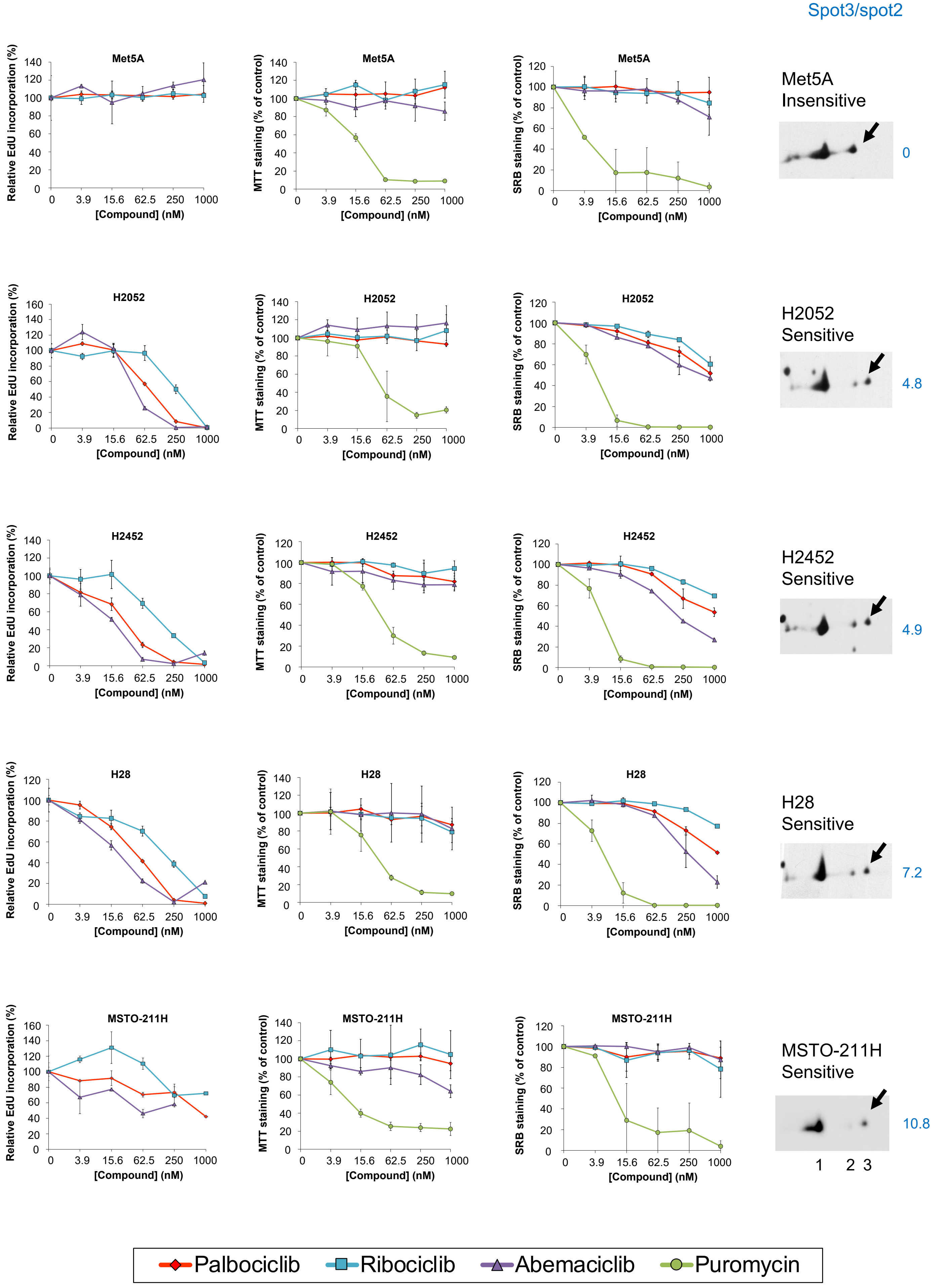

### Supplementary Figure S1B

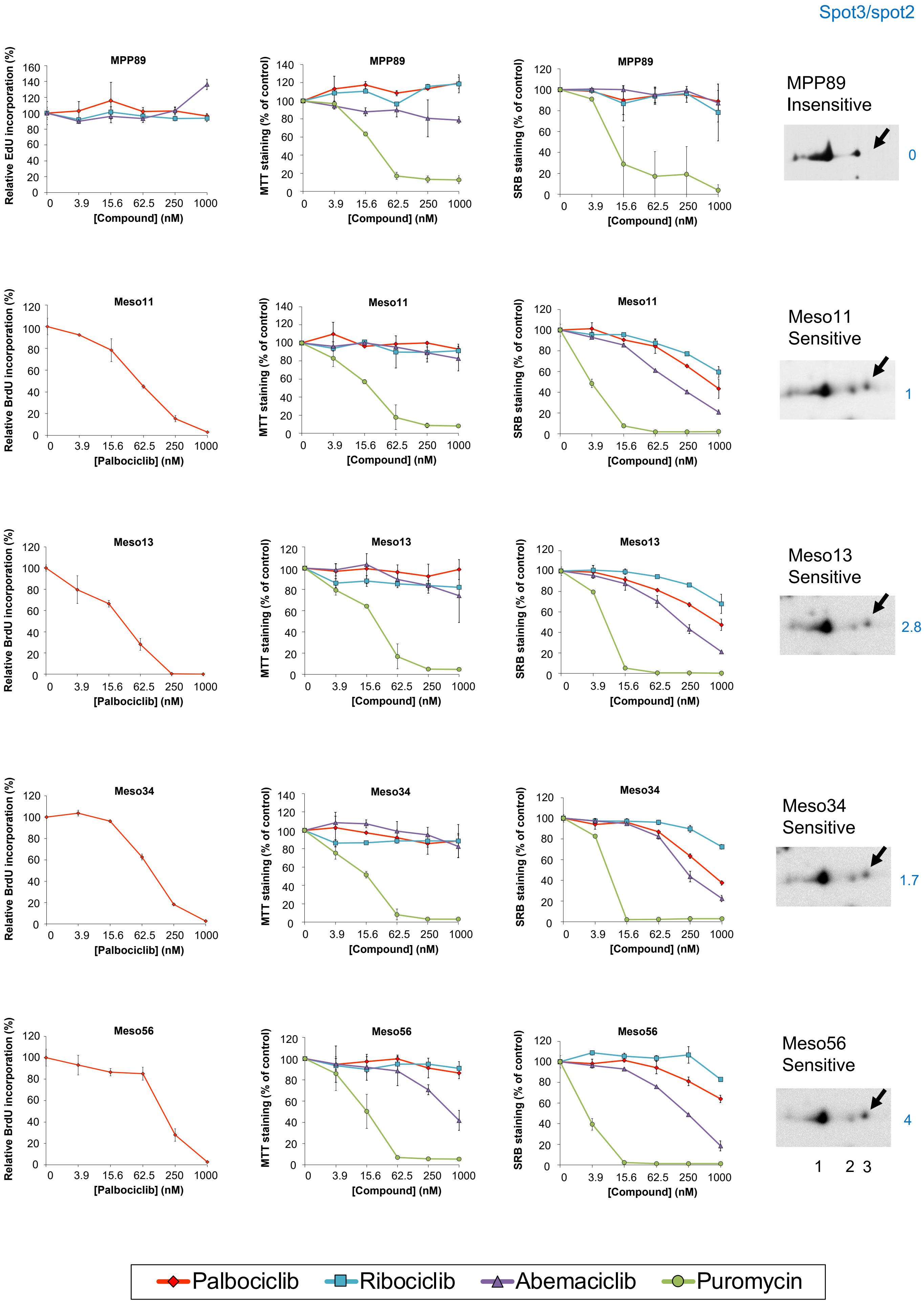

### Supplementary Figure S1C

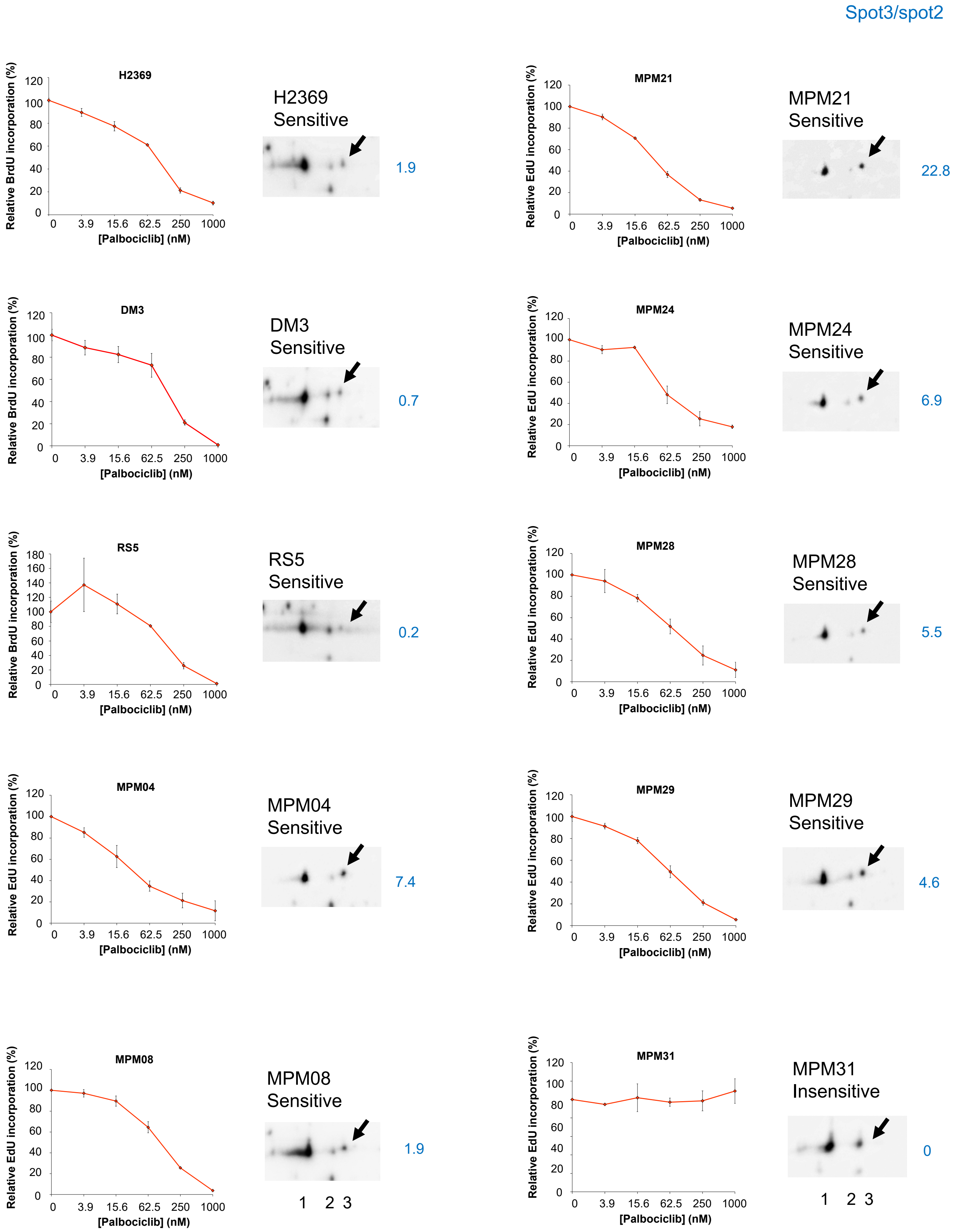

### Supplementary Figure S1D

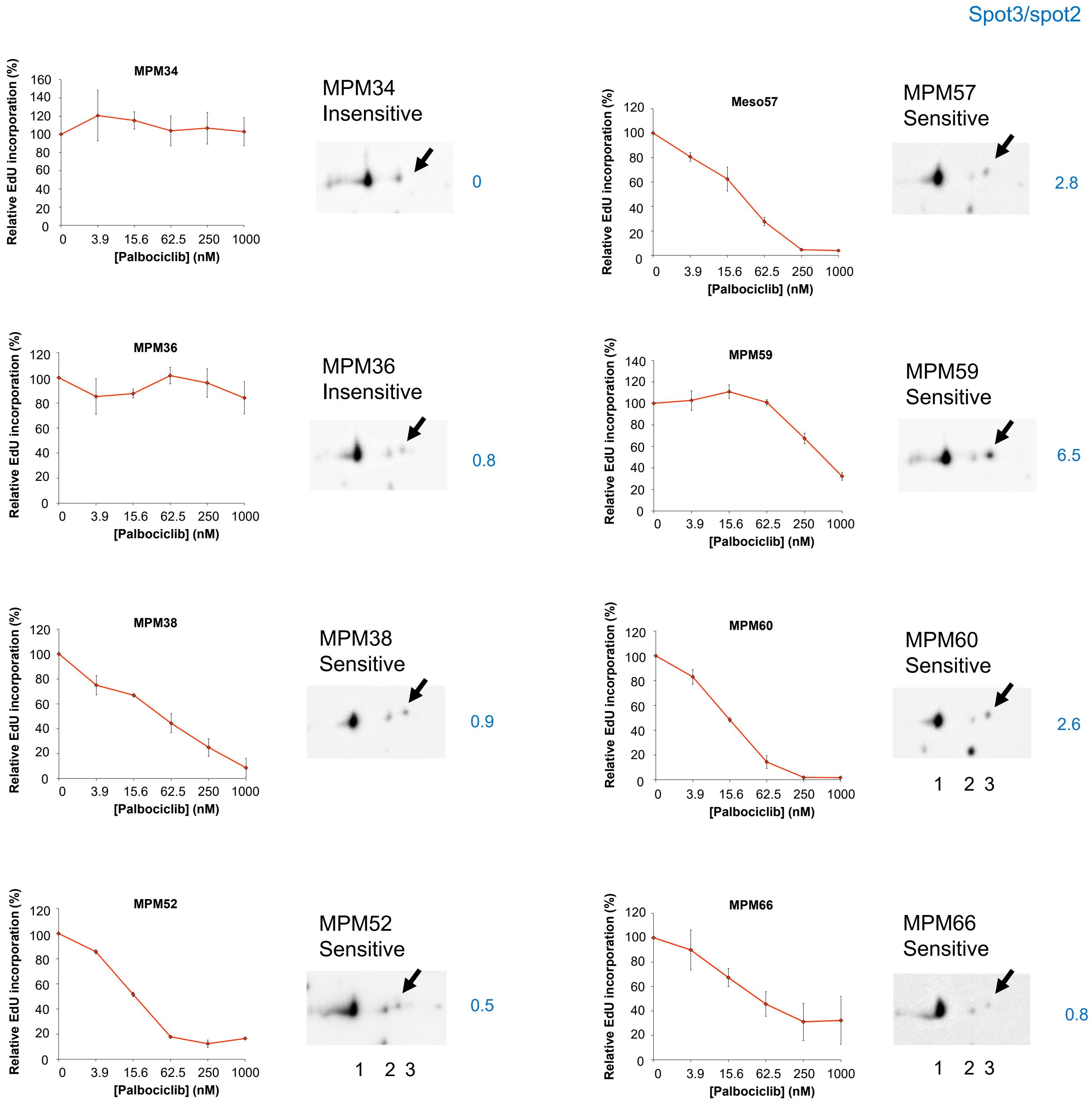

**Supplementary Figure S1.**
**Sensitivity of MPM cell lines to palbociclib is associated with CDK4 phosphorylation.**

Asynchronously growing MPM cell lines were treated for 24 h with DMSO, palbociclib, ribociclib or abemaciclib at the indicated concentrations. **A-D** DNA synthesis was evaluated from duplicated/triplicated dishes by counting the percentage of nuclei having incorporated BrdU/EdU during a pulse labeling of 1h. RS-5 cells were so slow growing that treatment with palbociclib was for 48 h with BrdU for the last 24 h. The relative proportion of BrdU/EdU-positive nuclei is expressed as percent of the mean value of control cells (**A-B** left panels, **C-D**). Error bars correspond to standard deviations scaled relative to the mean value of control cells except for the “MPM” cell lines (MPM_04 to MPM_66) where data represent mean +/- SD of 2 independent experiments. **A-B** The effect of CDK4/6 inhibitors was also measured after 48 h with MTT assay (middle panels) and after 6 days with the sulforhodamine assay (right panels). Increasing concentrations of puromycin were used as a positive control of cytotoxicity. Data represent mean +/- SD of 2 independent experiments. The profile of CDK4 separated by 2D-gel electrophoresis is also shown for each cell line. Arrows indicate the position of the T172 phosphorylated form of CDK4 (spot 3). The ratio of spot3/spot2 is displayed on the right of each panel.

**A** MeT-5A, H2052, H2452, H28, MSTO-211H

**B** MPP89, Meso11, Meso13, Meso34, Meso56

**C** H2369, DM-3, RS-5, MPM_04, MPM_08, MPM_21, MPM_24, MPM_28, MPM_29, MPM_31

**D** MPM_34, MPM_36, MPM_38, MPM_52, MPM_57, MPM_59, MPM_60, MPM_66

Supplementary Figure S2

**A**

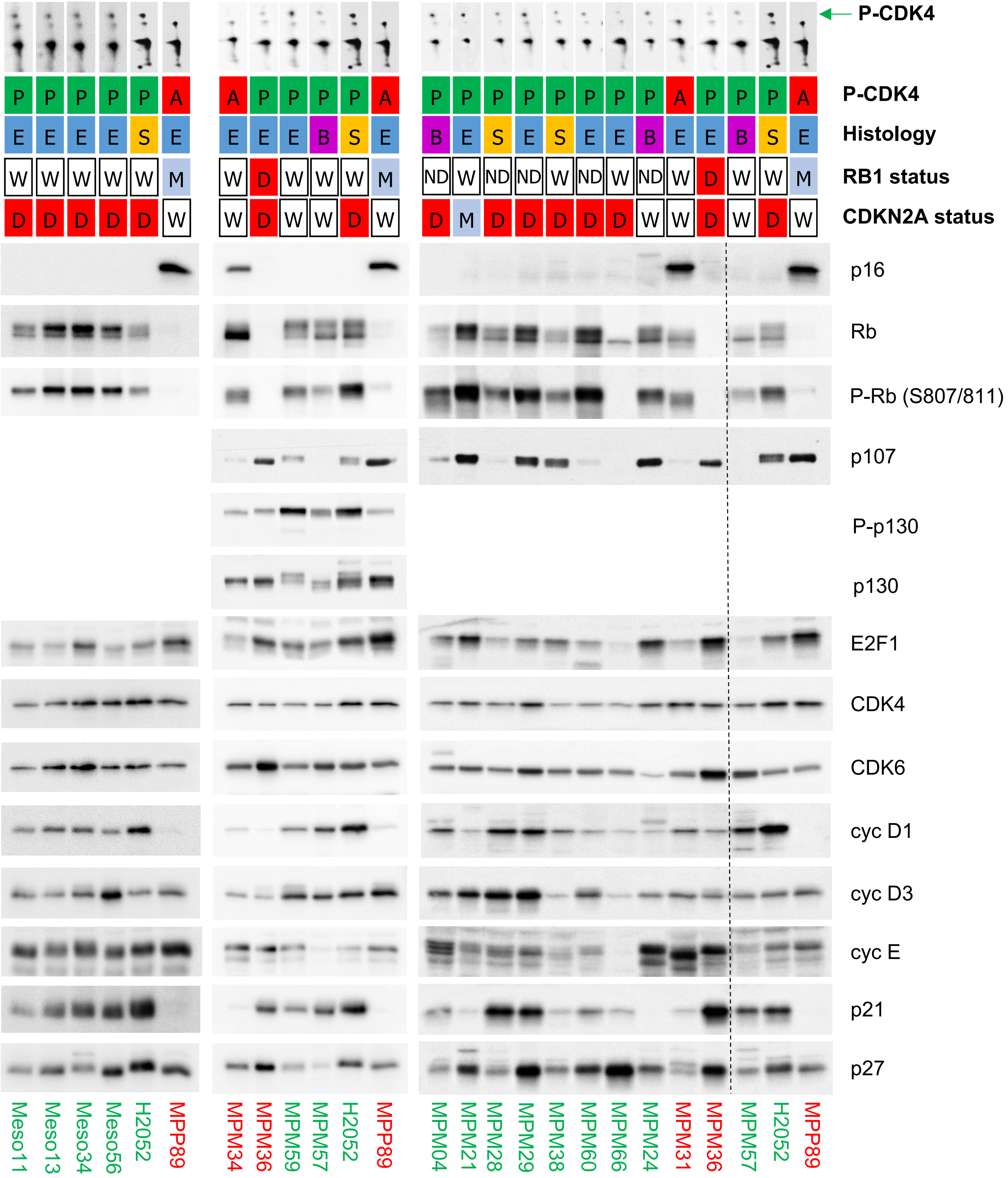


**B**

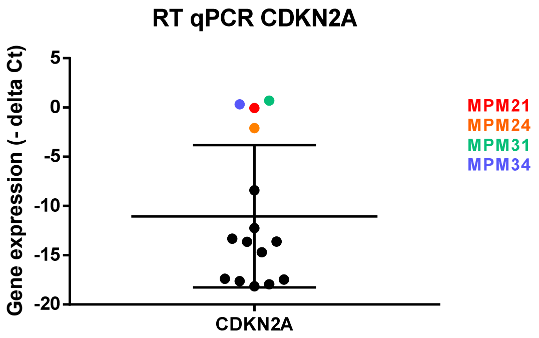

**Supplementary Figure S2.**
**Absence of CDK4 phosphorylation in resistant cells is due to high p16 levels associated or not with pRb defect.**

Sensitive cell lines are in green, resistant cell lines are in red.

**A** Total proteins were resolved by SDS-PAGE and detected with the indicated antibodies or separated by 2D gel electrophoresis followed by CDK4 immunodetection (the green arrow indicates the position of the phosphorylated form of CDK4). The following data are given for each cell lines: CDK4 phosphorylation profile (A: absent, P: phosphorylated), histology subtype (E: epithelioid, S: sarcomatoid, B: biphasic, N: normal), status of *RB1* and *CDKN2A* genomic locus (W: wild-type, D: deleted, M: mutated).

**B** mRNA expression of *CDKN2A* measured by RT-qPCR in MPM_04 to MPM_66 cells.

### Supplementary Figure S3

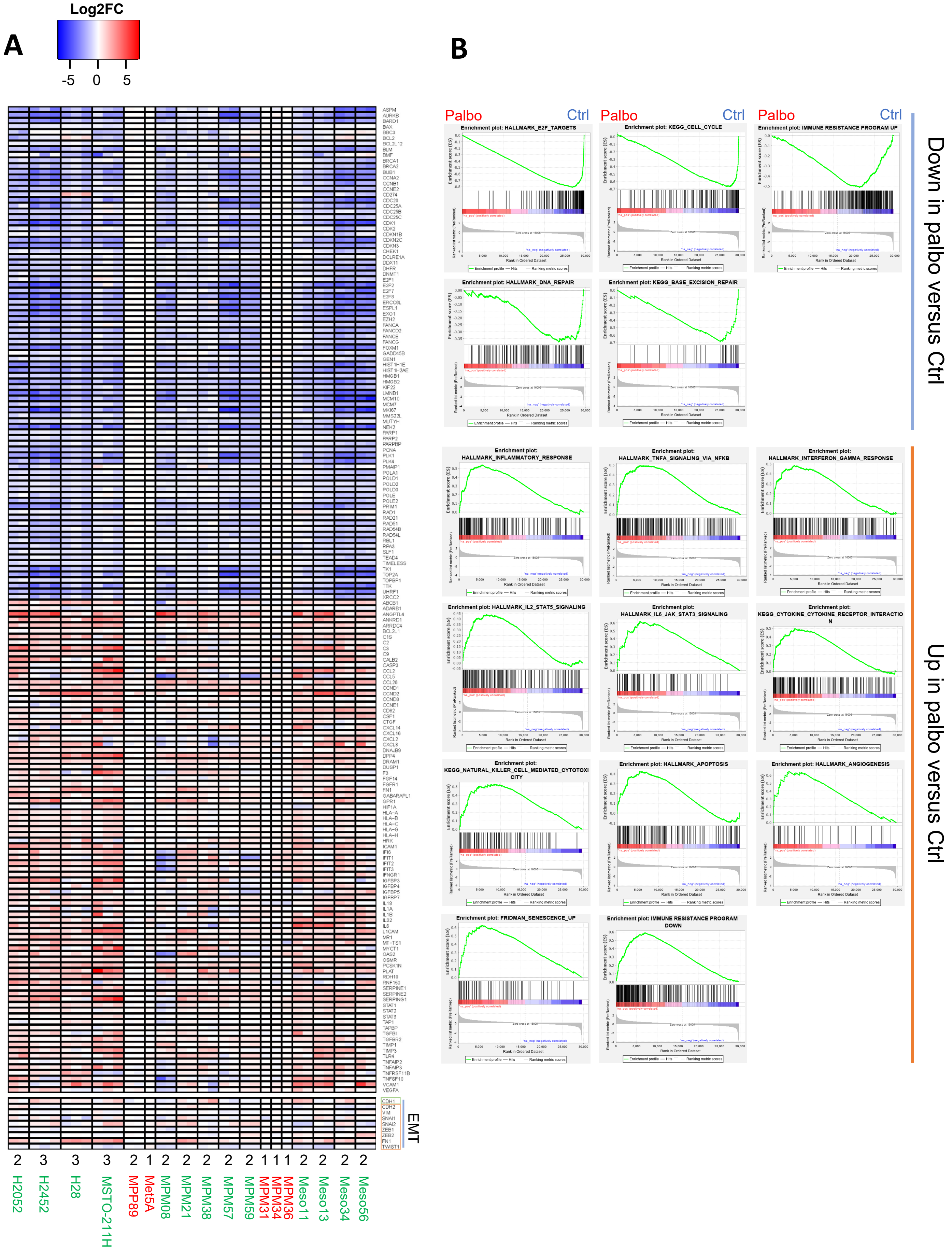

#### Supplementary Figure S3.
#### Impact of long term treatment with palbociclib analyzed by RNA-seq.

**A** Heatmap representing genes down- or up-regulated after a treatment of 9-10 days with 1 µM palbociclib. Genes involved in epithelial-mesenchymal transition (EMT) are grouped at the bottom of the plot (green frame: epithelial marker, orange frame: mesenchymal markers). Data are presented as log2FC (palbo/ctrl). n independent experiments are shown, as stated below the heatmap. Sensitive cell lines are in green, resistant ones are in red.

**B** GSEA enrichment plots for selected pathways down- or up-regulated by 1 µM palbociclib relative to control in a global analysis using all sensitive cell lines (n=13).

### Supplementary Figure S4

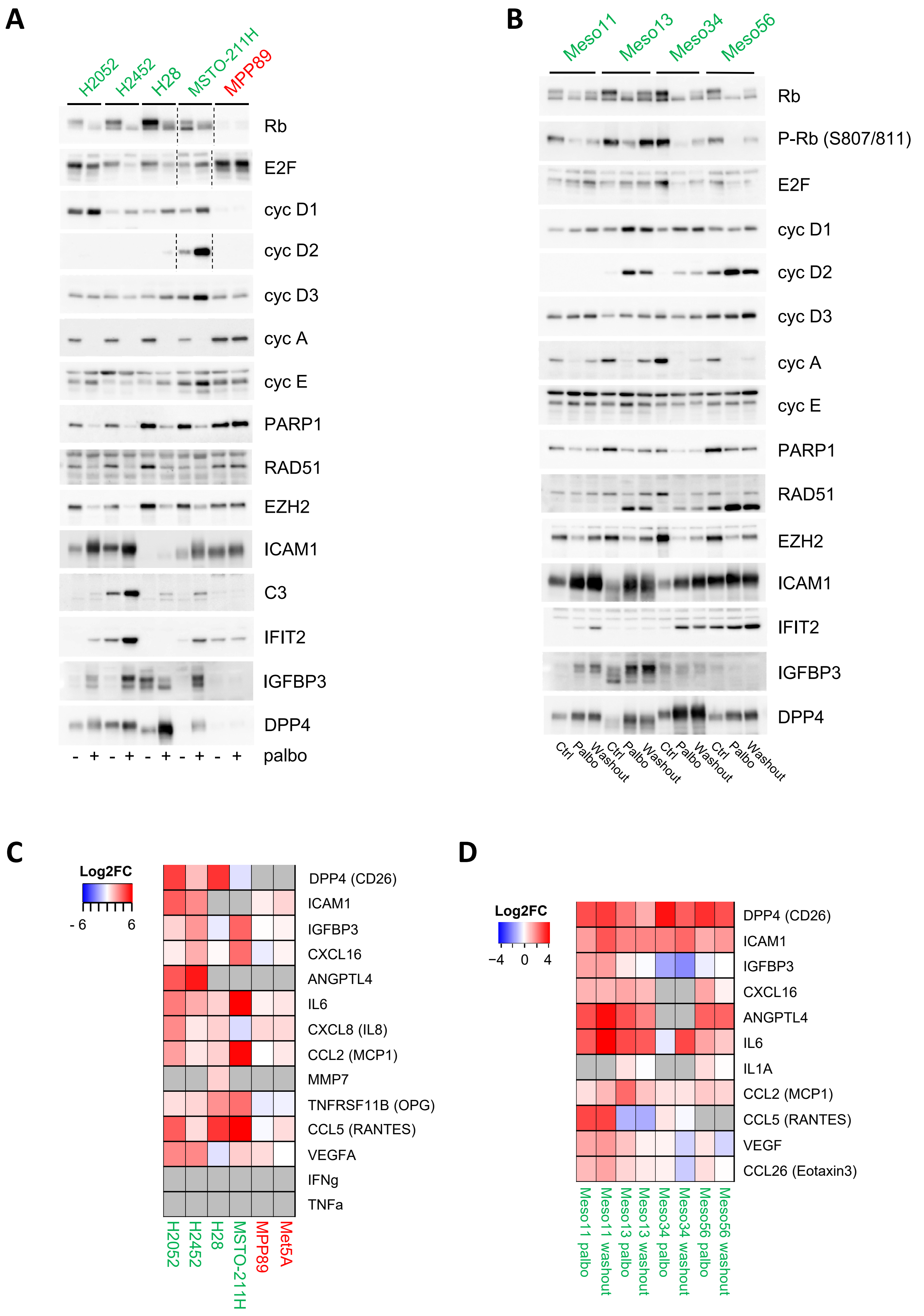

**Supplementary Figure S4.**
**Validation of RNA-seq data at the protein level**

Cells were treated for 9-10 days with DMSO (Ctrl) or palbociclib (Palbo) at 1 µM. In **B** and **D**, a drug washout of 48 h was also performed. Sensitive cell lines are in green, resistant ones are in red.

**A-B** Western Blot analysis with the indicated antibodies.

**C-D** Quantification of the indicated secreted proteins from cell culture supernatants by multiplex Elisa array. Conditioned media were collected 72 h (**C**) or 48 h (**D**) after the last change of medium. Heatmap represents log2FC (palbo/ctrl and washout/ctrl). Values under the detection limit are in grey.

### Supplementary Figure S5A

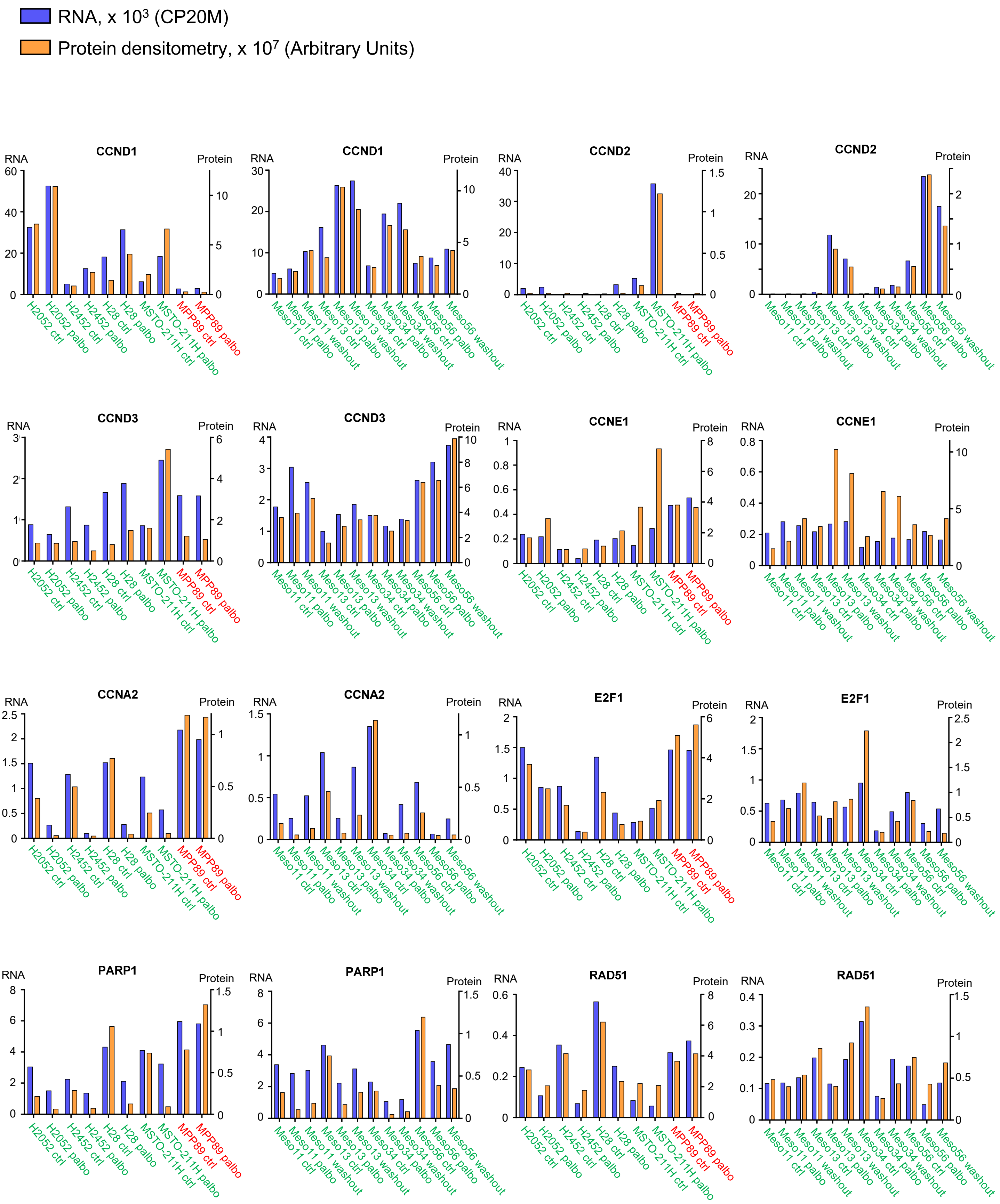

### Supplementary Figure S5B

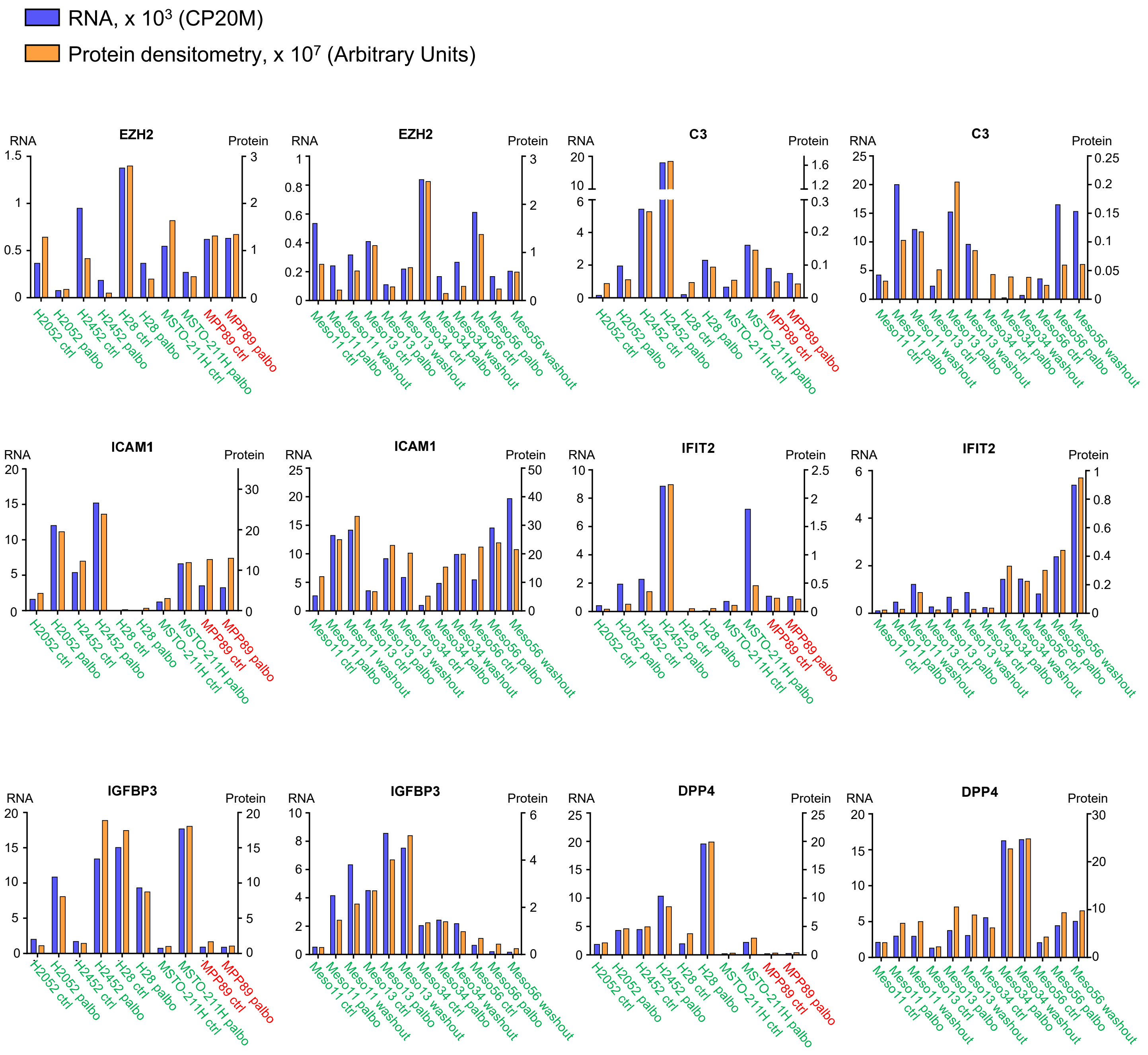

### Supplementary Figure S5C

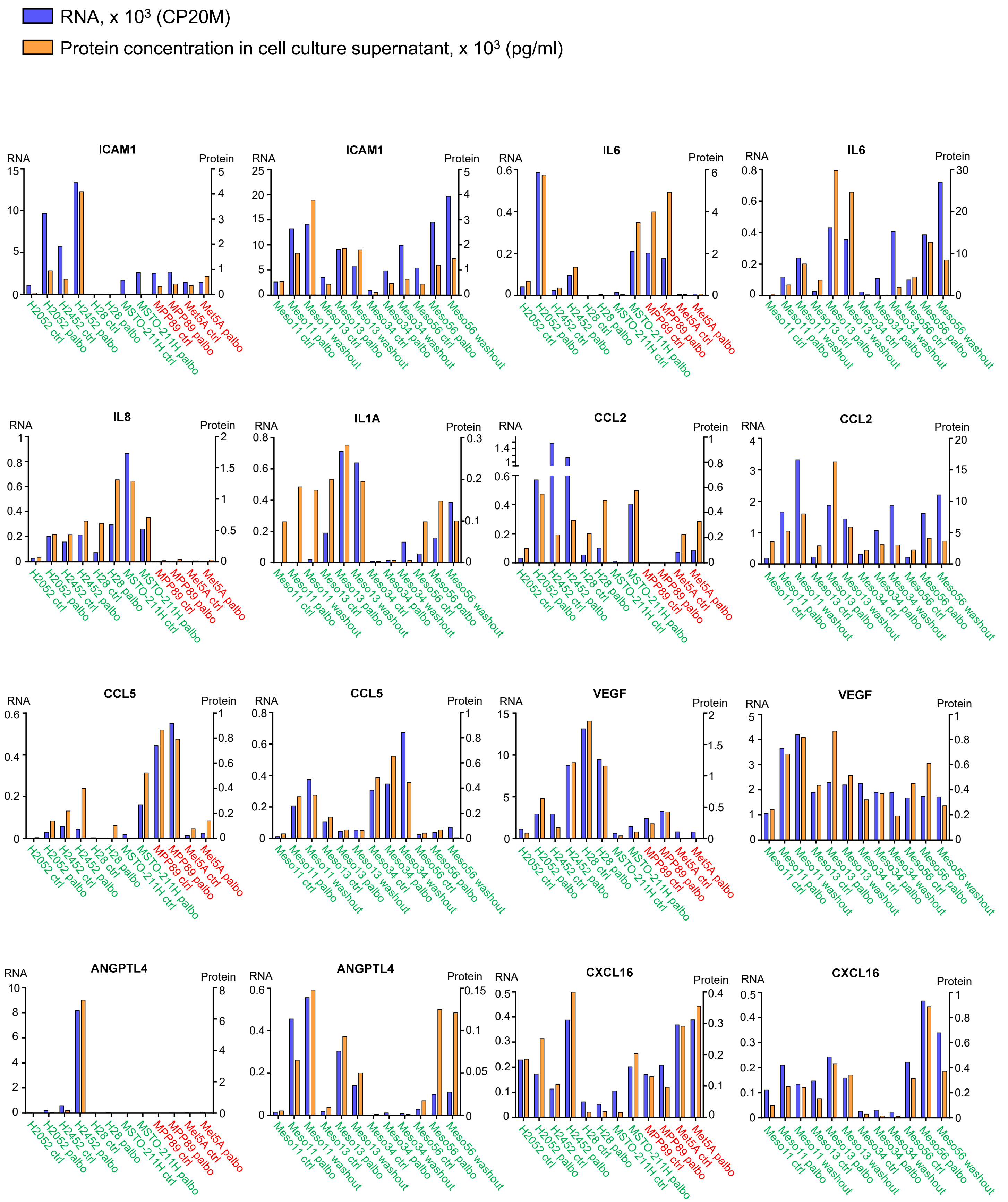

### Supplementary Figure S5D

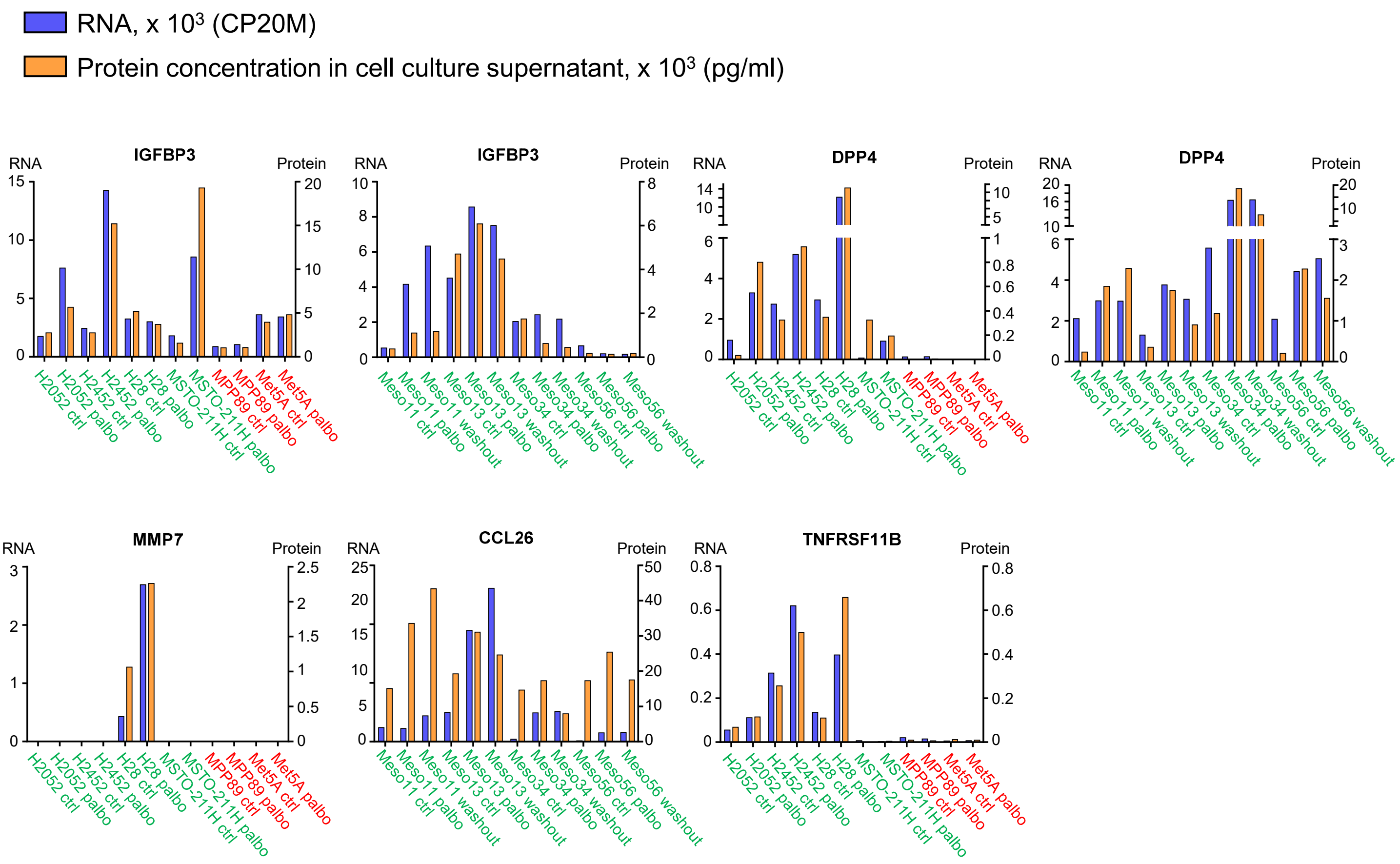


**Supplementary Figure S5.**
**Concordance between RNA-seq data and proteic analyses**

Sensitive cell lines are in green, resistant cell lines are in red.

**A-B** Plots combining, for indicated genes, mRNA levels from RNA-seq data normalized to the library size (CP20M) and the quantification of the corresponding protein from immunodetections.

**C-D** Plots combining, for indicated genes, mRNA levels from normalized RNA-seq data (CP20M) and the quantification of the corresponding protein from culture supernatants by multiplex Elisa.

### Supplementary Figure S6

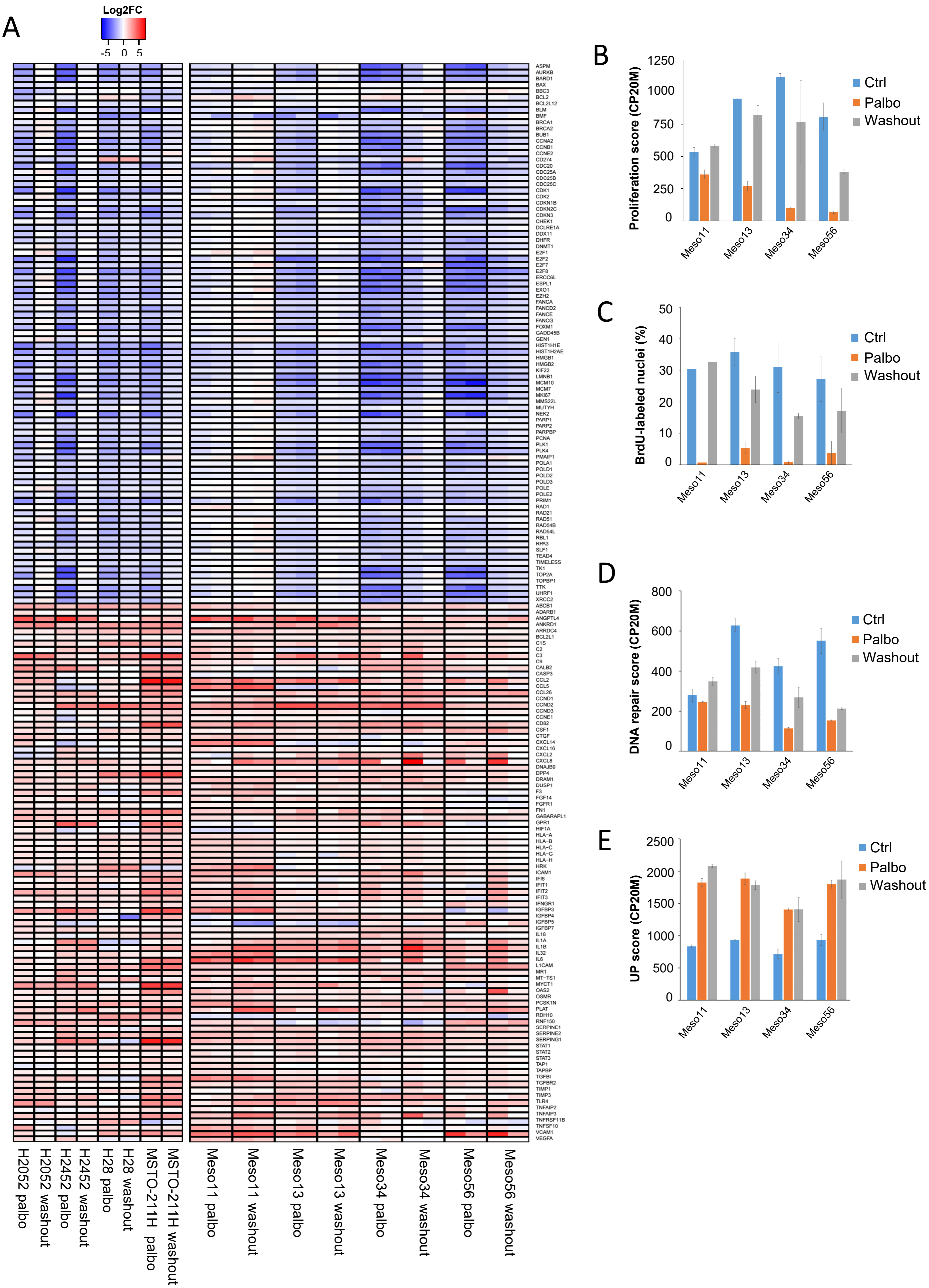

**Supplementary Figure S6.**
**Effect of palbociclib washout**

**A** Cells were treated for 10 days with 1 µM palbociclib followed or not by a drug washout of 48 h. Heatmap representing genes down- or up-regulated by palbociclib and the impact of drug washout on their expression. Data are presented as log2FC (palbo/ctrl and washout/ctrl). n independent experiments are shown, as stated below the heatmap.

**B** Proliferation score calculated from the RNA-seq data illustrated in **A** and expressed in counts per 20 million reads (CP20M). Mean +/- SEM of 2 independent experiments.

**C** DNA synthesis in cells treated as in **A**. Mean +/- SEM of 3 independent experiments except for meso11 (n=1).

**D** DNA repair score calculated from the RNA-seq data illustrated in **A** and expressed in counts per 20 million reads (CP20M). This score represents the median expression of the genes involved in DNA repair illustrated in Figure 5B. Mean +/- SEM of 2 independent experiments.

**E** Expression score of up-regulated genes calculated from the RNA-seq data illustrated in **A** and expressed in counts per 20 million reads (CP20M). Mean +/- SEM of 2 independent experiments.

### Supplementary Figure S7

Spot3/spot2
CDK4 modification profile

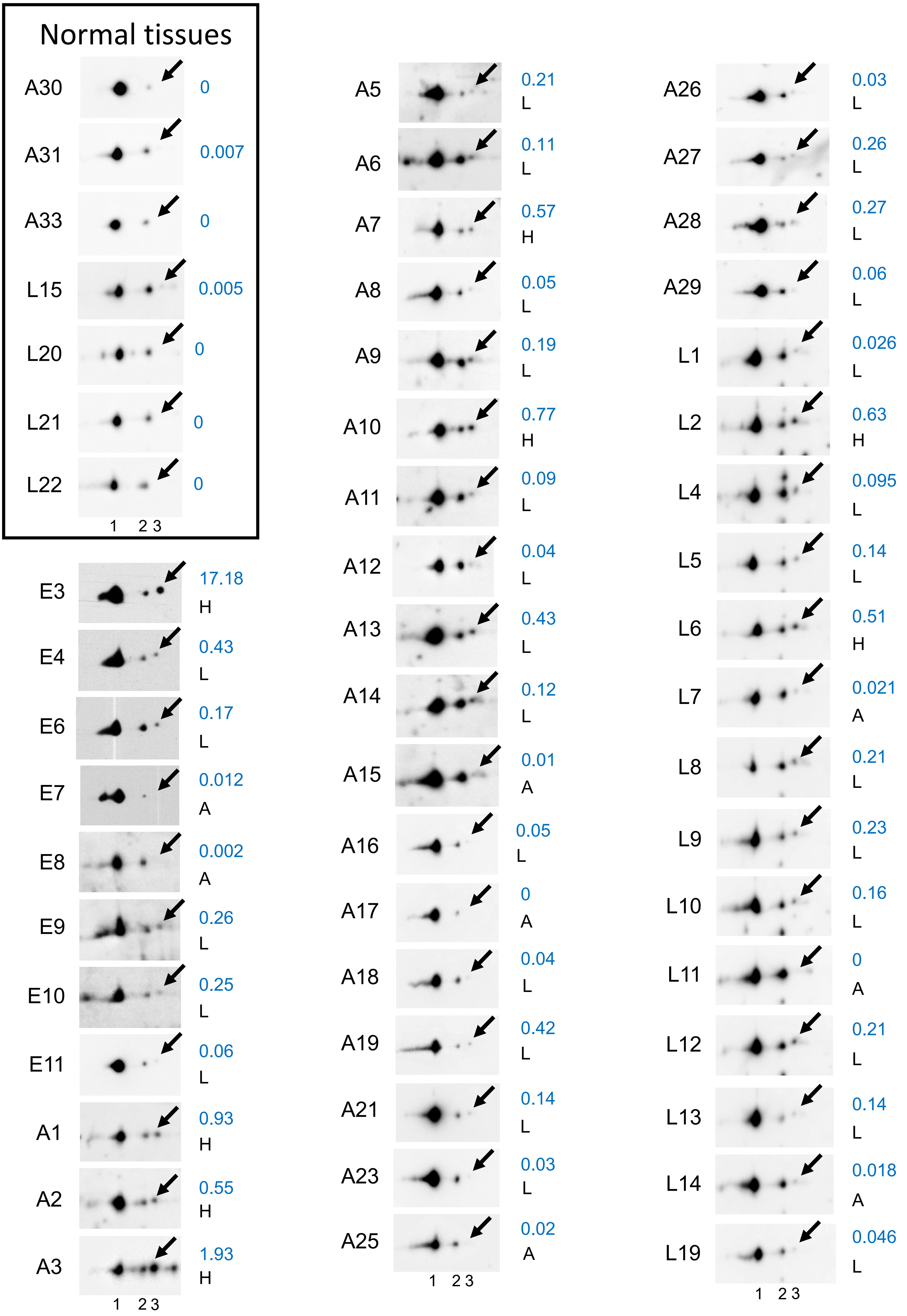

**Supplementary Figure S7.**
**CDK4 phosphorylation is present in most MPM tumors**

CDK4 immunodetection after separation by 2D-gel electrophoresis from 7 normal pleurae and 47 mesotheliomas. Arrows indicate the position of the T172 phosphorylated form of CDK4 (spot 3). Quantification of phosphorylated CDK4 (spot3/spot2) is indicated on the right and the CDK4 modification profile (H, L or A) is indicated for each tumor.

### Supplementary Figure S8

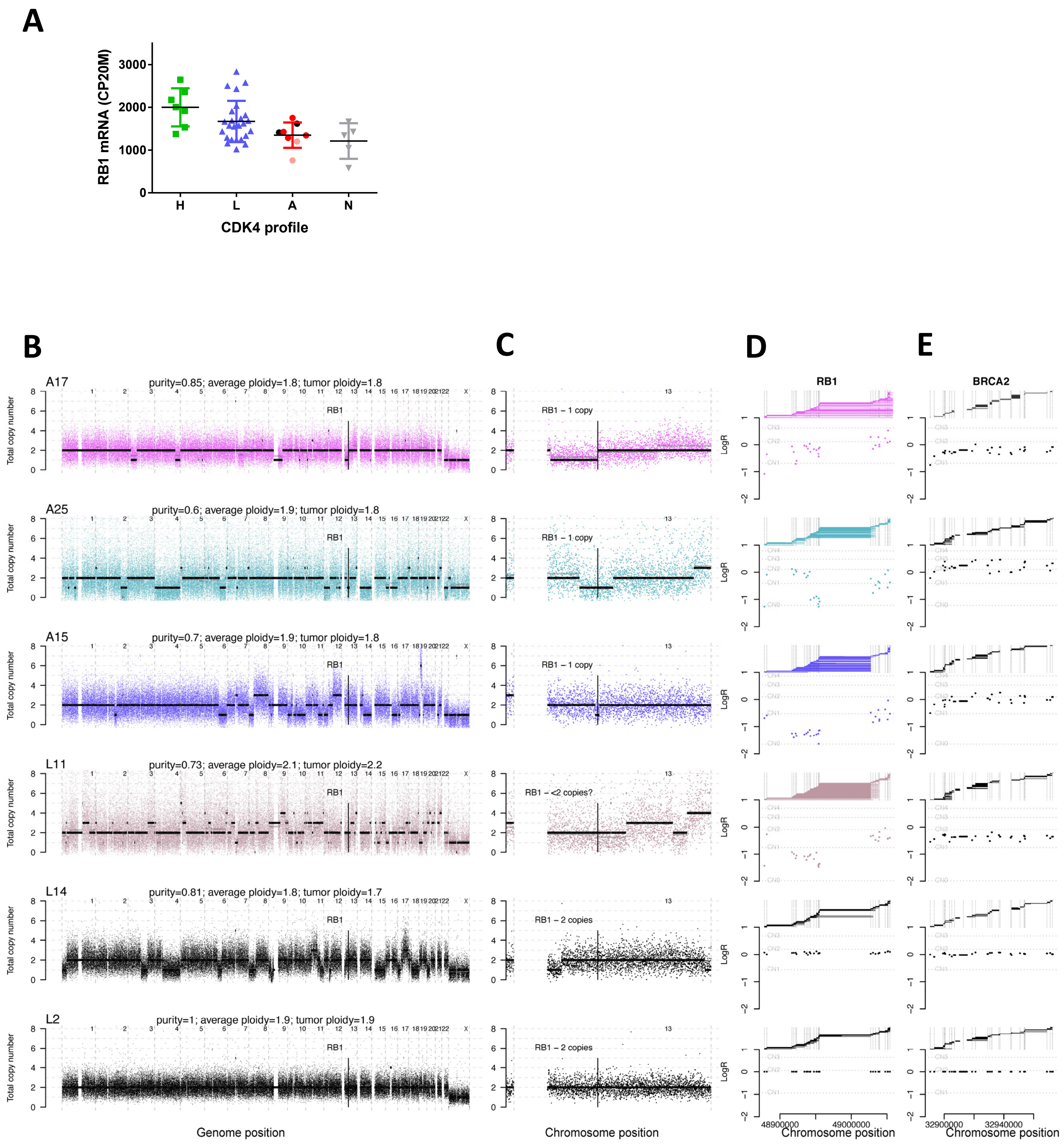

#### Supplementary Figure S8.
#### Double hits in *RB1* in high p16 expressing tumors: loss-of-heterozygosity and aberrant splicing.

**A** *RB1* mRNA levels (RNA-seq data in CP20M) in the different classes of tumors and in normal tissues. For class A tumors: red dots = absence of CDK4 phosphorylation with high *CDKN2A* expression and high proliferation score, pink dots = absence of CDK4 phosphorylation with low *CDKN2A* expression and high proliferation score, black dots = absence of CDK4 phosphorylation with intermediate *CDKN2A* expression and low proliferation score. Error bars: mean +/- SD. No significant difference between groups was found using the Kruskal-Wallis test.

**B-E** DNA-sequencing was performed for the four class A tumors with high p16 expression (A17, A25, A15 and L11). L14 and L2 tumors were used as controls.

**B** Genome-wide total copy-number profile across chromosomes. The raw underlying data is shown for each 30kb bins, the raw segment average (in gray) and the fitted rounded integer value (in black). The position of *RB1* is also shown.

**C** Zoom in chromosome 13 showing the number of copies of *RB1*.

**D** On the bottom graph, we show the logged read-depth ratios of the exon targets of *RB1* in the targeted DNA-sequencing data (y-axis) mapped to their genomic coordinates (x-axis). Horizontal lines show where the logged read-depth ratios would fall for different numbers of DNA copies, estimated from the sample purity and ploidy (Methods). On top of this graph, for each RNA-sequenced read that is mapped to an exon junction and sorted by its mapping position, we link it to its mate position. Horizontal lines therefore correspond to the junctions between exons like in a Sashimi plot. Vertical dotted lines indicate the position of the exons.

**E** Same as in **D** but for a control gene *BRCA2*.
